# Assessing Variations in Areal Organization for the Intrinsic Brain: From Fingerprints to Reliability

**DOI:** 10.1101/035790

**Authors:** Ting Xu, Alexander Opitz, R. Cameron Craddock, Margaret Wright, Xi-Nian Zuo, Michael P. Milham

**Author notes:** **Corresponding authors:** Michael P. Milham, M.D., Ph.D. Center for the Developing Brain, Child Mind Institute 445 Park Avenue, New York, NY 10022.

## Abstract

Resting state fMRI (R-fMRI) is a powerful in-vivo tool for examining the functional architecture of the human brain. Recent studies have demonstrated the ability to characterize transitions between functionally distinct cortical areas through the mapping of gradients in intrinsic functional connectivity (iFC) profiles. To date, this novel approach has primarily been applied to iFC profiles averaged across groups of individuals, or in one case, a single individual scanned multiple times. Here, we used a publically available R-fMRI dataset, in which 30 healthy participants were scanned 10 times (10 minutes per session), to investigate differences in full-brain transition profiles (i.e., gradient maps, edge maps) across individuals, and their reliability. 10-minute R-fMRI scans were sufficient to achieve high accuracies in efforts to “fingerprint” individuals based upon full-brain transition profiles. Regarding testretest reliability, the image-wise intraclass correlation coefficient (ICC) was moderate, and vertex-level ICC varied depending on region; larger durations of data yielded higher reliability scores universally. Initial application of gradient-based methodologies to a recently published dataset obtained from twins suggested inter-individual variation in areal profiles might have genetic and familial origins. Overall, these results illustrate the utility of gradient-based iFC approaches for studying inter-individual variation in brain function.

## INTRODUCTION

The delineation of functional brain units, commonly referred to as cortical areas (Van Essen and Glasser 2014), is a fundamental challenge in neuroscience. Central to such efforts, is the determination of areas of transition from one cortical area to the next. Pioneering efforts have used a combination of histological, cytoarchitectural, and myeloarchitectural examinations (Brodmann 1909; Vogt and Vogt 1919; Felleman and Van Essen 1991; Zilles and Amunts 2010), along with lesion studies, to differentiate neighboring cortical territories with respect to their architectonic, connectivity, functional and topographic properties (Felleman and Van Essen 1991; Cohen et al. 2008). Converging evidence supports the notion that boundaries between areas are generally sharp rather than gradual (Kaas 1987; Amunts and Zilles 2012). A well-defined example is the primary visual area (V1), which is clearly different and sharply separated in cell bodies from the second visual cortex (V2) (Sincich and Horton 2005; Buckner and Yeo 2014). Given that these measurements are mainly based on non-human primates, postmortem examinations, or invasive recording experiments, the development of methods capable of mapping cortical areas in vivo remains an active area of research.

Cohen et al. (2008) demonstrated the ability to map transitions between nearby cortical areas through the detection of spatial variation (i.e., gradients) in intrinsic functional connectivity (iFC) profiles estimated from resting-state fMRI (R-fMRI). A number of successful applications have since emerged. At the local level, studies have delineated transition zones between cortical areas within key regions of interest (e.g., cingulate cortex, left lateral parietal cortex, and inferior frontal cortex) (Cohen et al. 2008; Nelson et al. 2010; Hirose et al. 2013; Hirose et al., 2012). At the whole brain level, Wig et al. (2014) demonstrated the feasibility of delineating the broader range of boundaries between cortical areas in a single analysis based upon their gradient properties. Overall, these various studies have consistently recapitulated fine-grained cortical boundaries (e.g. between V1 and V2) previously established using histological and cytoarchitectural methodologies (Buckner and Yeo 2014). Additionally, recent studies demonstrated group-level transition-zone patterns to be highly reproducible across independent samples and studies (Gordon et al. 2014; Wig et al. 2014), thereby increasing enthusiasm towards the approach.

However, with few exceptions (e.g., Cohen et al. 2008; Wig et al. 2013; Laumann et al. 2015), studies focusing on detecting transition zones and cortical area boundaries have primarily relied on data averaged across dozens of individuals, or pooled individual-level maps to achieve more robust findings (e.g., Yeo et al. 2011; Kelly et al. 2012; Wig et al. 2014). Although effective in reducing noise, this is problematic, as a growing literature has suggested the presence of meaningful inter-individual variation in iFC patterns (Mueller et al. 2013), which appears to be stable over time and can be related to differences in behavior (e.g., Finn et al. 2015). Individual differences in areal organization are thought to be particularly prominent in higher-order association areas – a finding that is consistent with models suggesting the evolutionary value of cortical expansion (Mueller et al. 2013). Previous studies have suggested the potential importance of inter-individual variation in the transition zones or boundaries between cortical areas detected using iFC, by relating it to differences in task-evoked activation (Mennes et al. 2010) as well as variation in behaviorally quantifiable traits, such as social reciprocity and personality (Di Martino et al. 2009; Adelstein et al. 2011).

Recent work has underscored the feasibility of using iFC-based approaches to capture areal topographies at the individual level. First, Gordon et al. (2015) characterized the areal topological features at the individual-level, demonstrating that the areal sizes and the position of transition zones varied across individuals. Second, Laumann et al. (2015) mapped functional areal boundaries throughout the brain in a single human subject using a highly sampled dataset (>900 min). The individual-level architecture was broadly reflective of the universal architecture revealed in group-level analyses (Biswal et al. 2010; Wig et al. 2014), though with distinct variations that were fine-grained. Importantly, the areal organization was highly repeatable with subsets of data in this single subject rich dataset, suggesting that the generation of such highly sampled datasets is not a prerequisite for appreciating interindividual variation.

Here, we provide a multifaceted evaluation of test-retest reliability for transition-zones properties revealed by iFC-based mapping approaches (i.e., the gradients measured, and the boundaries drawn between functional areas delineated using them). To accomplish this, we estimated functional gradients and boundaries (i.e., edge maps) for each of 300 publicly available R-fMRI datasets (duration = 10 minutes; standard EPI sequence; TR = 2000ms), which were obtained from 30 participants who were scanned 10 times within one month. First we examined the “fingerprinting” potential of transition profiles (i.e., the ability to distinguish individuals from one another based upon their transition profiles). This was accomplished by testing whether the spatial correlation between any two sessions from the same individual was consistently greater than that observed between participants. Second, evaluated testretest reliability for indices of areal organization at the full-brain and vertex levels, using image intraclass correlation coefficient (I2C2) and intraclass correlation coefficient (ICC) respectively. For each of these perspectives, we repeated our analyses using 10-, 20-, 30-, 40- and 50 minutes to establish data needs to achieve moderate to high test-retest reliability. To ensure the generalizability of our findings, we replicate our analyses in the publicly available eNKI-TRT dataset, which made use of a multiband R-fMRI sequence (TR=645ms, duration=10 min). To provide a demonstration of the potential biological determinants of inter-individual variation in R-fMRI derived indices of areal organization, we demonstrated potential familial or genetic associations using a recently published R-fMRI dataset (Yang et al., 2016; Sinclair et al., 2015; TR = 2100ms, duration=5 min) obtained from 136 pairs of twins.

Of note, while initial work has based the determination of gradients and boundaries based upon a summary index for iFC, a multitude of alternative indices could be used. As such, additional analyses in the present work explore the similarities and distinctions among findings obtained using alternative iFC indices for the definition of gradients. Finally, we validated that the transitions and boundaries are not derived by the underlying structural architecture by comparing the results from real data to those from random data.

## METHODS

### Data Acquisition

Primary analyses in the present study were carried out sing the Hangzhou Normal University (HNU) test-retest dataset made publicly available via the Consortium for Reliability and Reproducibility (CoRR: http://fcon_1000.projects.nitrc.org/indi/CoRR/html/data_citation.html) (Zuo et al. 2014). The dataset consists of 300 R-fMRI scans, collected from 30 healthy participants (15 males, age = 24 ± 2.41 years) who were each scanned every three days for a month (10 sessions per individual) (Chen et al. 2015). Data were acquired at the Center for Cognition and Brain Disorders at Hangzhou Normal University using a GE MR750 3 Tesla scanner (GE Medical Systems, Waukesha, WI). Each 10-minute R-fMRI scan was acquired using a T2*-weighted echo-planar imaging sequence optimized for blood oxygenation level dependent (BOLD) contrast (EPI, TR = 2000 ms, TE = 30 ms, flip angle = 90°, acquisition matrix = 64 × 64, field of view = 220 × 220 mm, in-plane resolution = 3.4 mm × 3.4 mm, 43 axial 3.4-mm thick slices). A high-resolution structural image was also acquired at each scanning session using a T1-weighted fast spoiled gradient echo sequence (FSPGR, TE = 3.1 ms, TR = 8.1 ms, TI = 450 ms, flip angle = 8°, field of view = 220 × 220 mm, resolution = 1 mm × 1 mm × 1 mm, 176 sagittal slices). Foam padding was used to minimize head motion. Participants were instructed to relax during the scan, remain still with eyes open, fixate on a ‘+’ symbol, stay awake and not think about anything in particular. After the scans, all participants were interviewed to confirm that none of them had fallen asleep. Data were acquired with informed consent and in accordance with ethical committee review.

Two additional collections were included for secondary analyses. First, the Enhanced NKI-Rockland Sample test-retest dataset (eNKI-TRT), which is also available via CoRR (Zuo et al. 2014). Second, a recently published twin dataset obtained as part of the Queensland Twin Imaging (QTIM) study (de Zubicaray et al. 2008). The data acquisition and image preprocessing have been described in previous studies (Nooner et al. 2012; Zuo et al. 2013; Yang et al. 2016) and thus are only briefly outlined here.

The eNKI-TRT sample consists of data obtained from 22 participants (16 males, age = 33 ± 12.24 years) who were each scanned twice (1 week apart) at the Nathan S. Kline Institute for Psychiatric Research (Nooner et al. 2012). Each session includes a 10-minute R-fMRI scan (multiband EPI, TE = 30 ms, TR = 645 ms, flip angle = 65°, acquisition matrix = 112 × 112, in-plane resolution = 3 mm × 3 mm, 64 axial 3-mm thick interleaved slices, number of measurements = 404, multiband factor = 4) and a structural MRI (MPRAGE, TE = 2.52 ms, TR = 1900 ms, TI = 900 ms, flip angle = 9°, resolution = 1 mm × 1 mm × 1 mm). Data acquisition was performed on a Siemens 3T scanner with a 32-channel head coil located at NKI.

The QTIM study consists of data obtained from 200 twin pairs, including 86 monozygotic (MZ) and 114 dizygotic (DZ) twin pairs using a 4 T Bruker Medspec scanner (Bruker Medical). For each participant, the data consisted of 150 volumes of T2*-weighted BOLD contrast (EPI, TR = 2100 ms, TE = 30 ms, flip angle = 90, acquisition matrix = 64 × 64, field of view = 230 mm, 36 axial 3.6-mm thick slices). Highresolution T1-weighted images were also acquired (MPRAGE, TE = 3.83 ms, TR = 2500 ms, TI = 1500 ms, flip angle = 15°, slice thickness = 0.9 mm, acquisition matrix = 256 × 256 × 256).

### Image Preprocessing

All data were processed using the Connectome Computation System (CCS) (https://github.com/zuoxinian/CCS), which provides a platform for multimodal image analysis by combining components of AFNI (Cox 1996), FSL (Smith et al. 2004; Jenkinson et al. 2012), FreeSurfer (Dale et al. 1999; Fischl, Sereno, and Dale 1999), and SPM (http://www.fil.ion.ucl.ac.uk/spm). The CCS includes various implementations of quality control, surface-based R-fMRI measures, reliability and reproducibility assessments (Xu et al. 2015). Below summarize the structural and functional steps of the analyses employed.

Structural MRI preprocessing included spatial noise removal by a non-local mean filtering operation (Xing et al. 2011; Zuo and Xing 2011), followed by brain extraction, tissue segmentation, and surface reconstruction, as implemented in the standard recon-all processing pipeline in FreeSurfer 5.1 (http://freesurfer.net/fswiki). Tissue segmentation resulted in gray matter (GM), white matter (WM) and cerebrospinal fluid (CSF) masks for each hemisphere. For each participant, triangular meshes representing white matter and pial surfaces were reconstructed by tessellating the GM-WM and GMCSF interfaces, and averaging the surfaces (white matter, pial) to create a middle cortical surface (Dale et al. 1999; Fischl, Sereno, and Dale 1999). The resulting surface in native space was then inflated into a sphere for alignment to the *fsaverage* template by shape-based spherical registration (Fischl, Sereno, Tootell, et al. 1999). Following calculation of the transform for registration to the fsaverage, which has 164,000 vertices, we then down-sampled the surface to a representation with 10,242 vertices (fsaverage5) to ensure computational feasibility of the functional analyses to be performed in the present work. Quality control was carried out using a series of screenshots generated by CCS to facilitate visual inspection of the outputs from essential processing procedures. In particular, for the preprocessing of a given structural MR image to be considered valid, the outputs of the following steps had to pass inspection: brain extraction, tissue segmentation, surface reconstruction, co-registration and normalization.

Functional preprocessing included: discarding the first five volumes of the R-fMRI time series and storing all images as double precision floating point numbers (to reduce variability of outcomes across operating systems, see Glatard et al. 2015), detecting and compressing temporal spikes (AFNI 3dDespike), slice timing correction (this was not performed for multiband images), motion correction (3DVolReg), and 4D intensity normalization to 10,000. Nuisance variable regression (Fox et al. 2005; Lund et al. 2006) was performed to remove the mean timeseries for individual-specific WM and CSF masks (derived from the FreeSurfer segmentation), as well as 24 motion parameters (Friston et al. 1996; Yan et al. 2013); linear and quadratic trends were also removed from the data. The time series residuals were bandpassed filtered (0.01-0.1 Hz) to restrict the signals to those frequencies previously implicated in resting state functional connectivity (Biswal et al. 1995; Cordes et al. 2001). Functional images were co-registered to the native high-resolution anatomical images using 6-parameter boundary-based registration (BBR) in FreeSurfer (Greve and Fischl 2009).

### Quality Control Procedure

Following the functional preprocessing, the frame-wise displacement (FD) and mean FD was calculated to quantify the head micromovements (Power et al. 2012, 2014; Patriat et al. 2013). Any scans with mean FD greater than 0.2 mm were excluded from our analysis. For the testretest HNU dataset employed in our primary analyses, mean-FD was less than 0.2 mm for all 300 scans (mean=0.057, SD=0.019). In addition, the percentage of excessive motion frames (FD >0.2 mm) for each of the scans was found to be less than 1%.

For the eNKI-TRT, 20 participants (age=33.40 [SD=12.58], 14 male) passed our criteria (mean FD < 0.2 mm) and were included in the current studies (mean=0.107, SD=0.036). For the QTIM twin dataset, 379 of 400 participants passed the head motion criterion (mean FD < 0.2 mm). Given that previous studies have raised concerns about the potential contributions from gender difference in brain structure (e.g. cortical area, thickness, gyrification, etc.) and head motion (Schmitt et al. 2007; Chen et al. 2013; McKay et al. 2014; Docherty et al. 2015) to findings in twin studies, Yang et al. (2016) included only same sex adult twin pairs in which both twins exhibited low motion (mean FD < 0.2mm). We used the same criteria and resulting participant list in the present work. This resulted in inclusion of 272 participants (age=22.12 [SD=2.35], 68 male), consisting of 78 MZ twin pairs and 58 DZ twin pairs in our analysis. These criteria controlled the potential heritability of the head motion and spatial information; no significant differences between MZ and DZ pairs were found in the mean FD (p=0.52); only a nonsignificant trend was noted between MZ and DZ pairs with respect to the level of spatial correlation for anatomical images in MNI space (p=0.08).

### Computation of iFC Metrics on Native Surface

There are several advantages for analyzing fMRI data on the surface rather than in volume space. A cortical sheet provides a more accurate representation of the morphology and topology of brain structure (Dale et al. 1999). It provides a more accurate registration between individual data and the template surface (Van Essen 2004; Ghosh et al. 2010; Klein et al. 2010; Yeo et al. 2010). Also, it enables the visualization of spatial relationships between brain regions in terms of their geodesic distances along the cortical surface, which is more neurobiologically meaningful than 3D Euclidean distance in volume space. In light of these putative benefits, all functional indices in the present study were calculated on the surface. First, the volumetric fMRI data were aligned to anatomical space using the BBR transformation matrix and then projected to their corresponding native middle cortical surface (about 160k vertices for each hemisphere). To make the resolution comparable between the volumetric and surface data, a coarser *fsaverage5* template surface was used (the average triangle edge length between adjacent vertices of fsaverage5 is about 3.5 mm) for computing functional metrics. The transformation calculated during spherical registration was applied to the *fsaverage5* template to convert it into a standard mesh of the native surface, for which each node has a direct correspondence with a node on the fsaverage5 surface (Van Essen, Glasser, Dierker, Harwell, et al. 2012). The standard mesh was calculated by interpolating the native surface to a transformed version of the fsaverage5 template in native space.

Consistent with previous studies of connectivity gradients, iFC similarity was measured by the spatial similarity between whole brain functional connectivity maps calculated from a seed vertex and those calculated from every other voxel’s time courses (Cohen et al. 2008; Wig et al. 2011, 2014). Specifically, for each individual, the time course for each vertex was extracted and used to calculate a whole brain functional connectivity profile that consists of 20,484 vertices (10,242 for each hemisphere) of cerebral grey matter and 9,413 voxels in subcortical regions and the cerebellum. The distributions of the resulting correlation values were standardized to the normal distribution using Fisher’s r-to-z transform. For each hemisphere, an iFC similarity map was measured for each vertex by calculating the spatial correlation between the vertex’s iFC profile and the iFC profile of every other vertex - resulting in a 10,242 vertices × 10,242 vertices symmetric matrix. Each column (or row) of this matrix represents the vectorized iFC similarity map for each surface vertex. These 10k iFC similarity maps were then smoothed along the native surface with a Gaussian Kernel (FWHM=8mm, about twice of the surface resolution [~3.5mm], see Supplementary Figure S1–2 to address the smooth effect). The gradient (i.e., the first spatial derivative) of each smoothed iFC similarity map was computed on the native middle surface to measure transitions in iFC profile across vertices, resulting in 10k gradient maps for each hemisphere. The details of the gradient computation are described in the next section.

Beyond iFC similarity, we adopted three additional functional indices that capture local-, global- and network-scale aspects of intrinsic brain function, respectively, and can each be used to generate gradient maps (independent of iFC similarity). Two-dimensional Regional Homogeneity (2dReHo) was employed as a local-level measure of iFC that characterizes the temporal synchronization of the BOLD signal at each vertex across the cortical mantle (Zang et al. 2004; Zuo et al. 2013; Jiang et al. 2014; Jiang and Zuo 2015). For each vertex, ReHo was defined by the Kendall’s coefficient of concordance of the time series within the given vertex and its one-step neighboring vertices. The preprocessed R-fMRI data for ReHo calculation was not spatially filtered, but temporally band-pass filtered to avoid artificial increases of ReHo intensity (Zuo et al., 2013).

The network-scale measure of iFC was captured using a dual regression (DR) procedure to map 10 intrinsic connectivity networks previously defined from a meta-analysis of activations from the BrainAtlas database (Smith et al. 2009). These networks includes Medial Visual network (DR-MedVis), Occipital Visual network (DR-OccVis), Lateral Visual network (DR-LatVis), Default Mode network (DR-DMN), Cerebellar network (DR-Cerebellar), Sensorimotor network (DR-SenMot), Auditory network (DR-Audi), Executive Control network (DR-Control), Left Fronto-parietal network (DR-FrontalL), and Right Fronto-parietal network (DR-FrontalR). The spatial network maps were conducted in the first regression on normalized R-fMRI data in MNI152 space to generate the characteristic time series of 10 networks for each individual. The resultant characteristic time series were further entered into the second regression to yield the spatial map of 10 networks on cortical surface in individual level.

At the full-brain scale, Degree centrality (DC) and Eigenvector centrality (EC) were employed to capture features of iFC as a whole (i.e., full-brain connectivity patterns) (Bullmore and Sporns 2009; Rubinov and Sporns 2010; Zuo et al. 2012). Specifically, pairwise iFC matrix was calculated for all possible pairings of vertices on the cortical surface using Pearson’s correlation. We then thresholded and binarized the full brain iFC matrix at *p*(r)<0.001.

### Transition Zones, Boundaries and Parcellation

Gradients were calculated on the convoluted (i.e., not flattened) standardized native surface mesh, followed by an edge detection procedure that sharpened the gradients to identify putative boundaries. The initial framework for gradient mapping follows the Canny edge detection procedure (Canny 1986) and involves three steps: (1) each vertex’s iFC connectivity profile is smoothed with a 2D Gaussian filter to reduce noise, (2) a gradient map is calculated for each vertex’s smoothed profile to identify regions of rapidly changing IFC profiles, and (3) performing edge detection by finding local maxima in the gradient maps. Cohen et al. (2008) first applied Canny edge detection to iFC similarity profiles calculated from data projected onto flattened surfaces. Flattening the cortical surface requires it to be cut in five places; findings associated with edges near these cuts are prone to artifact and are thus difficult to interpret. To avoid this issue, Wig et al. (2013, 2014) extended the method to unflattened surfaces by performing all three steps (smoothing, gradient computation, edge detection) on the middle cortical surface (mid-way between white matter and pial surfaces). The major challenge of this approach is that the gradient calculation must account for the curvature of the cortical surface. This was handled by calculating the gradient of each vertex on a plane perpendicular to the surface normal vector extending from that vertex (Caret 5.65 ‘metric-gradient’ function http://brainvis.wustl.edu/wiki/index.php/Caret:About). The gradient is calculated by solving a linear model relating the geodesic distances between the vertex and it’s nearest neighbors to differences in their image intensity. The magnitude of the resulting gradient is the change of image intensity per unit distance. Here, the intensity image is a functional feature of the cortical surface described in the above section, including iFC similarity, ReHo, DC, EC and DR-Networks. The gradients were calculated for each of these functional indices to detect abrupt changes at the local, global, and network scales.

Finally, the parcels and corresponding edges were identified from the resulting gradient maps using a “watershed by flooding” algorithm (Gordon et al., 2014). This algorithm identifies the local minima on a gradient map as starter seeds for parcel creation, and then gradually grows the seeds outward until they meet another parcel. The barriers built in this procedure for separating parcels are identified as edges. With the exception of iFC similarity, an individual-specific binary edge map was generated for each gradient map (ReHo, DC, EC, and DR-networks).

For the iFC similarity map, a more complex set of procedures is carried out, generating an edge density map for each individual, rather than a simple binary edge map. Specifically, for each vertex, a gradient map was calculated, as well as a corresponding edge map using the watershed algorithm - resulting in 10k gradient/edge maps for each hemisphere. For each individual, the 10k gradient/edge maps were then averaged to generate final gradient map and an edge density map. While an edge density map can be more preferable depending on the specific application, it is also possible to generate a binary map by applying the same watershed algorithm to the edge density map.

To make the resultant maps from ReHo, DC, EC and DR-networks more comparable to those iFC similarity, we created edge density maps for these R-fMRI measures by bootstrapping. Specifically, we generated alternative observations of the data using circular block bootstrap to ensure the temporal dependencies of R-fMRI data (Bellec et al. 2008; 2010). The procedure concatenated temporal blocks of original data (block length = the square root of the number of total time points) to form a time series with same temporal points as the original. Each temporal block was randomly extracted from the original time series with replacement. We utilized 100 bootstraps to yield 100 edge maps. The final edge density maps were calculated by averaging 100 bootstrapped binary edge maps for ReHo, DC, EC and DR-networks.

For each of the R-fMRI measures (iFC similiarity, ReHo, DC, EC, DR-networks), the resulting discrete parcels were assigned to large-scale system networks driven using group-averaged parcellation generated by Gordon et al. (2015). The assignment procedures are described in the next section.

### Network Systems Assignment

Recent studies have demonstrated that the network structures of parcellation in individuals were similar to the group averaged system (Gordon et al., 2015; Laumann et al., 2015). To characterize large-scale systems for each edge map generated in the present work, we assigned system identity to parcels created in the above procedure using the previously established network definitions from Gordon’s (2014). The matching procedure was modified based on Gordon et al., (2015). Specifically, we first detected the vertex with local minimal edge density for each discrete parcels. Then we averaged that time series with the time series of its one-step neighbors within that parcel. The averaged time series was considered as a representative time series of the chosen parcel, and correlated against all other time series across cortical surface to obtain a connectivity map. After that, we thresholded and binarized the connectivity map at the top 5% of connectivity strengths. This resulted in a binarized map of regions with high connectivity to that local minimal vertex. Then we measured the overlap of this binarized map to the 12 network maps from Gordon, the best match network with highest Dice similarity was assigned to that vertex and its representative parcels. Consistent with prior work, the neighboring parcels with the same network identifier were merged together to create a “network-patch”.

### Estimating Intra-, Inter-Individual Variability and Intra-Class Correlation

Intra-and inter-individual variability were simultaneously estimated using separate Linear Mixed Effects Models (LMEs) for gradient and edge density maps. The gradient or edge density for a given vertex *v* can be denoted as *Y*_*ij*_(*ν*), where *i* indicates the participant and *j* represents the measurement (for *i*=1, 2,…, 30 and *j*=1, 2).

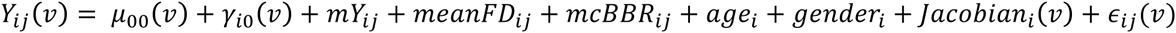

In this model, *μ*_00_(*v*) is a constant term that represents the intercept or fixed effect of the group average in gradient/edges at vertex *v*, while γ_*i*0_(*ν*) is the random effect term for i-th participant at vertex *v*. Due to the potential influence of various confounding factors to our final estimation, we included several covariates in this mixed model at both the session and individual level. At the session level *j*, the model included three covariates: the mean frame-wise displacement (*meanFD*_*ij*_) of head motion, minimum cost of BBR (*mcBBR*_*ij*_), and the global mean of gradient/edge across the whole brain within group mask (*mY*_*ij*_) for each participant *i*. At the individual level, age_*i*_, gender_*i*_ and Jacobian determinant of the spherical transformation (*Jacobian*_*i*_*ν*) at vertex *v*) were included for each participant *i* as well. The model estimations were implemented using the *Ime* function from the *nlme* R package (http://cran.r-project.org/web/packages/nlme). The variance estimation 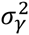 of the random effect term *y*_*i*0_(*ν*) is the inter-individual variance across all the participants, while the variance 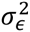 of the residual *ϵ*_*ij*_ is the intraindividual variance for a single participant across all the sessions. Meanwhile, the intra-class correlation (ICC) was calculated by dividing inter-individual variance to the sum of inter-and intra-individual variance.

### Surface Geometry Constraints Testing

To determine whether transition zones and detected boundaries could have arisen from the surface geometry, we replicated the boundary mapping analyses using surrogate data. Specifically, for each participant, we replaced the preprocessed R-fMRI data with temporal Gaussian random data (mean = 0, standard deviation = 1, the same number of time points as the original data) in native functional volume space. The surrogate data were then projected onto the participant’s native middle surface, registered and down resampled to ‘*fsaverage5*’ version of native surface using the same co-registration and spherical registration transformation calculated during processing of the real data. The functional measures (iFC similarity) were computed, followed by gradient computation and edge detection using the same procedure and registration transformations used for the real data. The gradient and boundaries maps from this surrogate data were compared with the results from real data to investigate whether surface geometry and registration errors would contribute to the functional areal organization detected.

## RESULTS

### Boundary Map Characteristics

For each of the 300 scans (30 participants, 10 sessions each) from the HNU data collection, a gradient map and a corresponding edge map were generated. This was accomplished using the gradient-based boundary mapping approach and iFC similarity measure specified by Wig et al. (2014), though at the individual level. Specifically, for each individual, the maps generated were averages of those calculated for each of the 20k vertices (10k per hemisphere). Prior to examining differences among participant-specific maps, individual level maps were averaged to generate a group-level gradient map that was strikingly similar to those previously published by Wig et al. (2014) and Gordon et al. (2014) (Figure 1A). To measure the similarity of our group map versus previous work, we calculated the percentile of the group-level gradient scores for the vertices that were identified as brain systems boundaries in Gordon et al. (2014). The median percentile of gradient within Gordon’s system boundaries of was 70.62%. The group-level edge density map was also similar to those previously published (the median percentile of edge density was 63.79% within Gordon’s boundaries), though somewhat less comparable; this was largely due to the fact that we calculated the edge maps on individual participants prior to averaging, as opposed to group-average maps (e.g., Gordon et al. calculated an edge map for each vertex using the group-level gradient map, and then averaged across edge maps). The latter approach averages out the contributions of individual variation in the gradients prior to the generation of the edge maps (see supplementary materials Figure S3 for a direct comparison of the two approaches from prior methodological testing by our group using the QTIM dataset.

**Figure 1.**
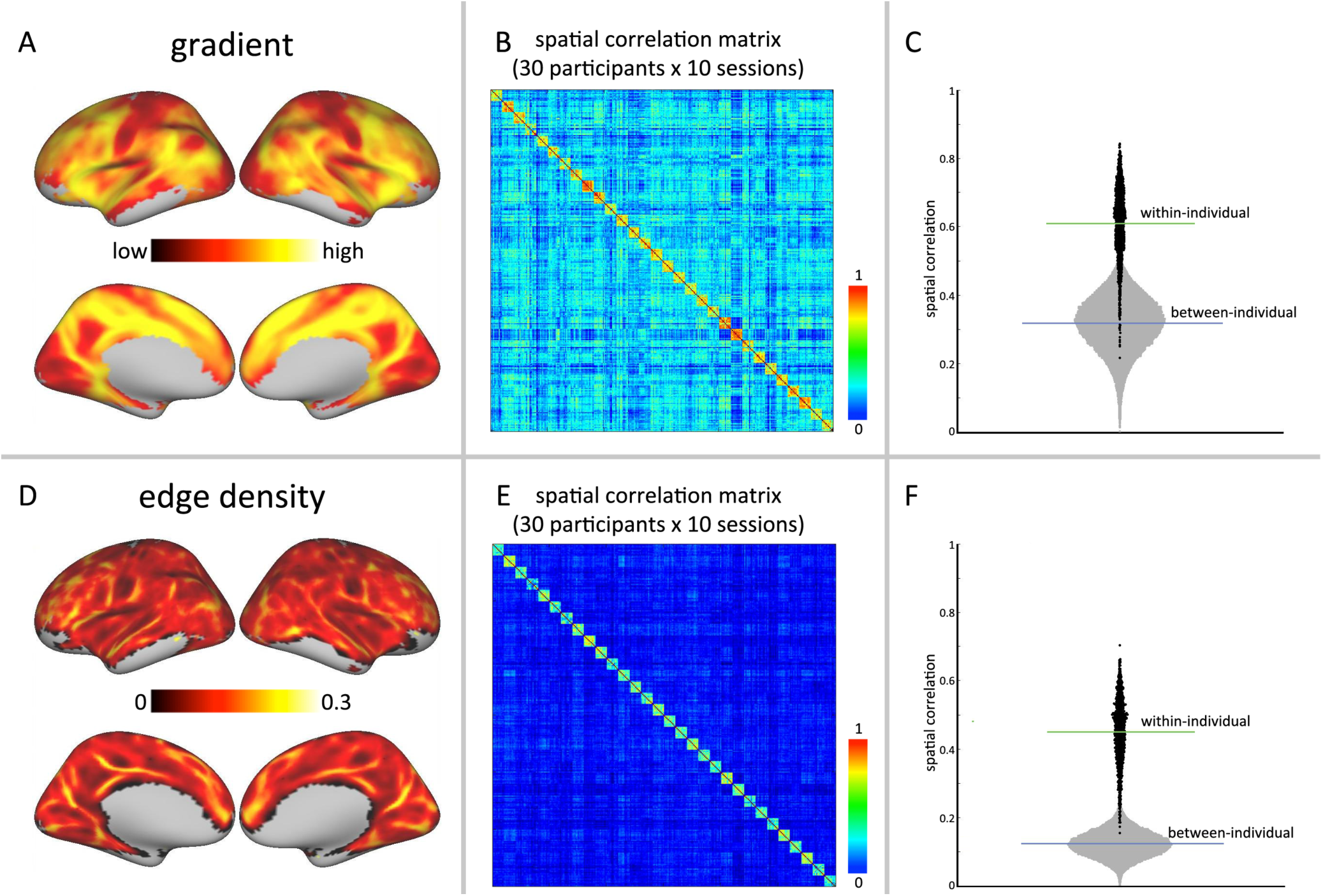
Individual areal organization is unique to each individual and highly repeatable across scan sessions. A: Group-level gradient and edge density maps for iFC similarity. The individual maps were calculated on the standardized *‘fsaverageö’* version of native surface for each individual, then averaged and visualized here on the *‘fsaverageö’* template surface. B: Between-scan spatial correlation matrices for the gradient (top) and edge density (bottom) maps derived from the 300 datasets (30 participants * 10 sessions); the rows/columns of the matrix are ordered by participant (i.e., first ten rows are sessions 1 to 10 for participant one, second ten rows are sessions 1 to 10 for participant two and so on; total number of participants = 30). C: Distribution of between-scan spatial correlations (from Figure 1B). Black dots are the within-individual (i.e., same participant, different sessions) correlations for gradient (top) and edge density maps (down), while the grey dots are between-individual correlations (i.e., different participants, same session, or different participants, different sessions).

### High Reproducibility of Individual Areal Boundaries (Fingerprinting)

For both gradient maps and edge maps, we next investigated the similarity of the maps generated across individuals and time (i.e., scan sessions). In order to accomplish this, we calculated the spatial correlation between the 300 gradient maps generated (30 participants * 10 sessions), as well as between the 300 edge maps. Figure 1B demonstrates the spatial correlation matrix obtained for gradients (upper panel) and edge maps (lower panel). Regardless of participant, the correlations between different scan sessions (gradient: mean r=0.609 (SD=0.110); edge: mean r=0.450 (SD=0.093)) are notably greater than those between participants (gradient: mean r=0.318 (SD=0.078); edge: mean r=0.123 (SD=0.037)). To facilitate appreciation of this point, the correlation matrix is ordered by participant (i.e., first ten rows belong to participant 1, second ten rows belong to participant two, and so on). As can be seen in Figure 1C, the distribution of between-individual correlation scores for gradient maps overlap minimally with that for within-individual; the correct identification rate was 90.44% (1221 in total 1350 within-individual correlations of 300 scans); For edge maps, fewer overlaps were present for within-and between-individual correlations, introducing 99.26% correctly identified rate. These findings suggest that the full-brain areal transition characteristics indexed by gradient and edge measures are unique to each individual, and repeatable across scan sessions.

Figure 2 depicts data from multiple 10-minute sessions for three representative participants to provide illustrative examples of distinct patterns associated with each participant, as well as their repeatability.

**Figure 2.**
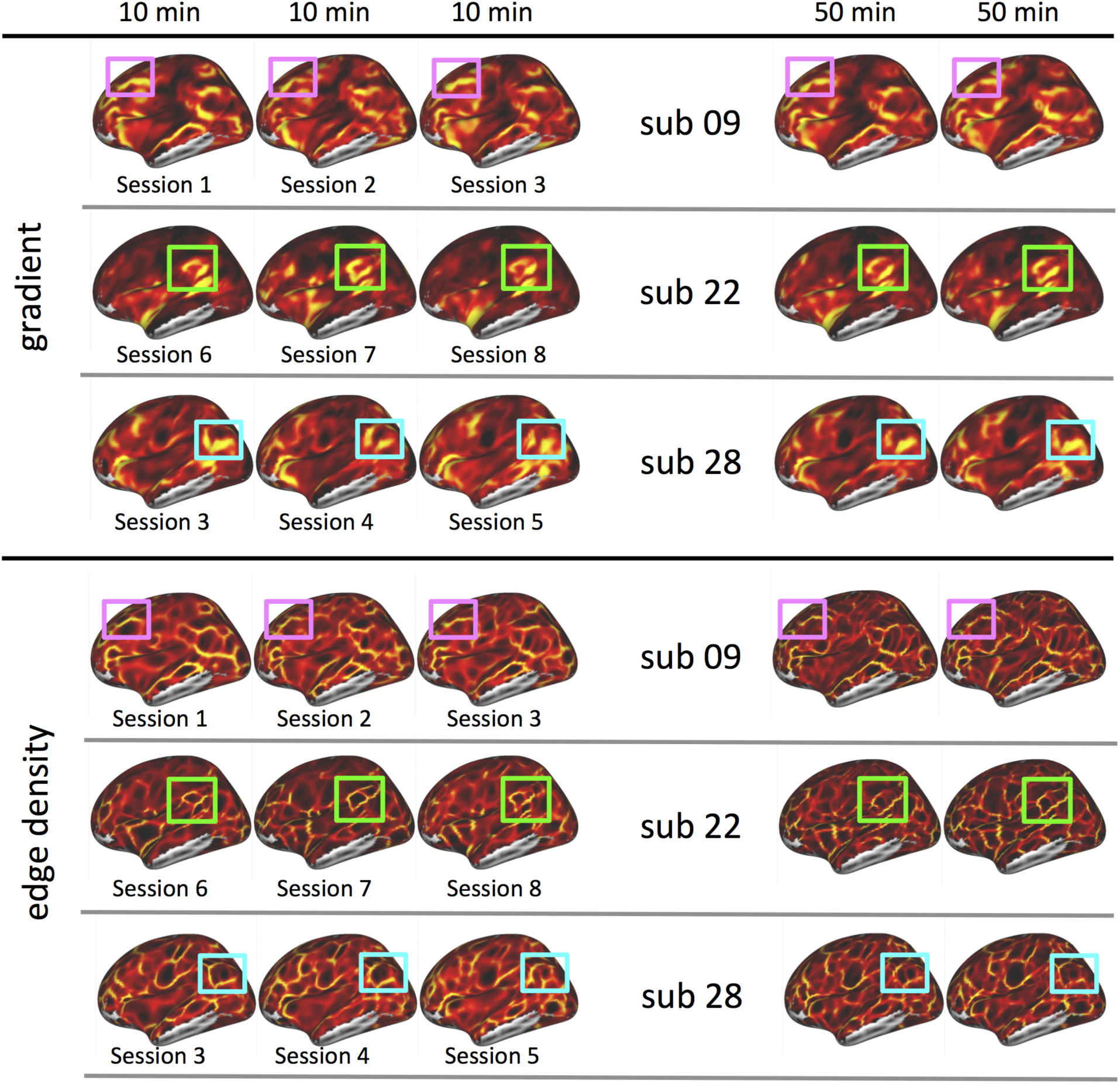
Examples of gradient and edge density of iFC similarity in 3 individual participants. Left three columns are gradients from three sessions and right two columns are gradients from two 50-minutes subsets. To facilitate comparison, for each participant we mark off (using colored boxes) an example of a feature that differs across participants and is relatively consistent within individual.

The distribution of within- and between- individual spatial correlations for each individual was plotted in Figure S4. We further examined whether the within-individual spatial correlations were related to age, gender, head motion or the functional-to-structural coregistration. We did not find any significant relationships between the mean or standard deviation of within-individual correlations and the factors listed above.

We replicated the general pattern of findings in another independent R-fMRI dataset (eNKI-TRT data collection: 20 participants * 2 sessions, multiband EPI sequence TR=645ms). Specifically, we again observed substantially higher within-individual correlations (gradient: mean r=0.56 (SD=0.159); edge: mean r=0.364 (SD=0.095)) than between-individual correlations (gradient: mean r = 0.219 (SD=0.129); edge: r=0.075 (SD=0.032)).

### High Reproducibility of Individual System Patches

Regarding to system-scale patch, we used Dice Similarity to measure the reproducibility of within- and between-individual scan sessions for the 12 network patches defined by Gordon et al. (2014). Five of the 12 network patches (visual, auditory, dorsal sensorimotor, ventral sensorimotor, default) exhibited relatively high reproducibility across both sessions and individuals (i.e., average pairwise Dice Similarity > 0.5) (see Figure 3) – suggesting a relatively conserved architecture for these networks.

**Figure 3.**
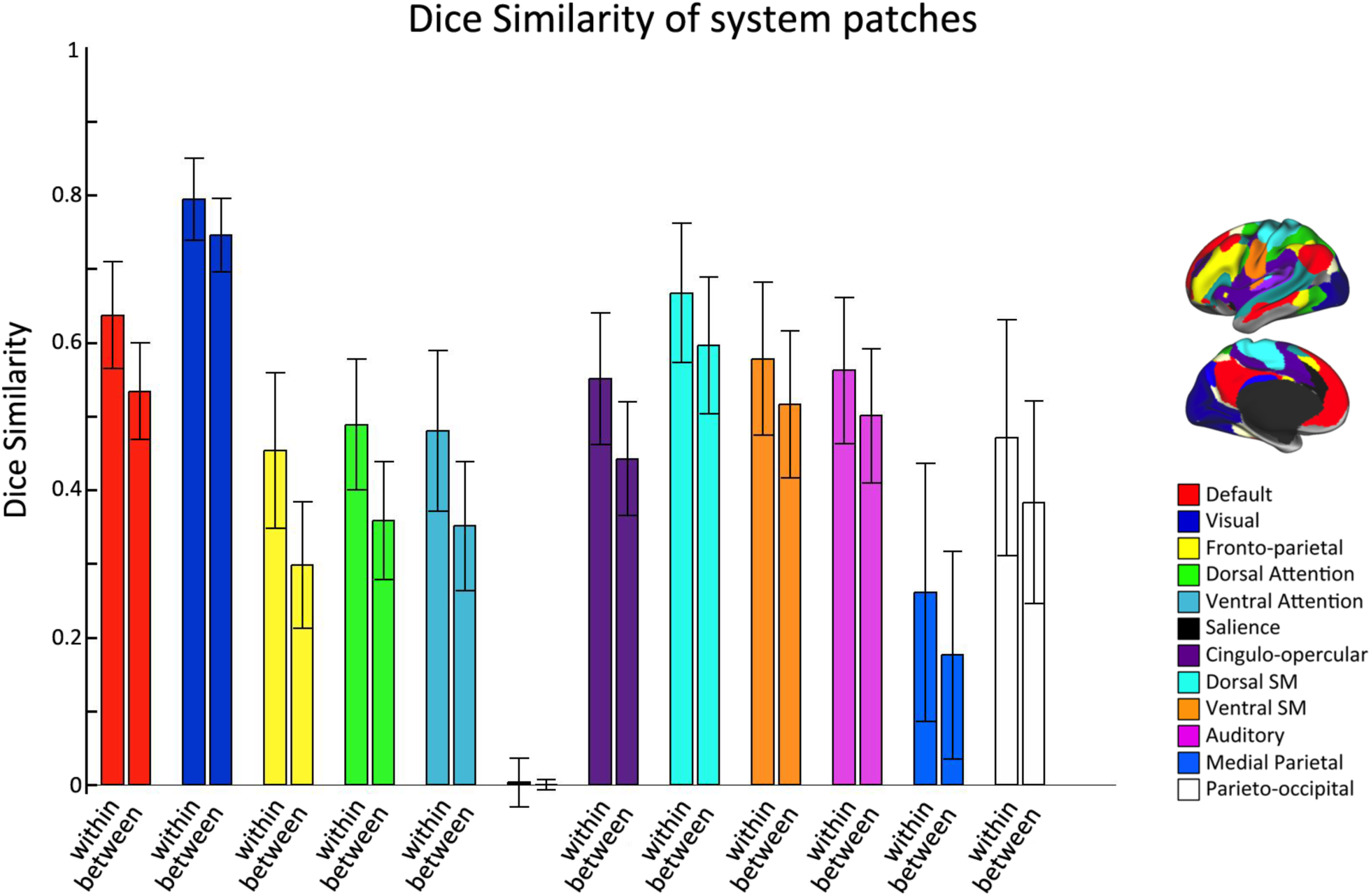
The reproducibility system-matched patches across 10 sessions and 30 participants. The color-coded bars indicate Dice similarity of system patches (mean and standard deviation) for within and between individual at each system network.

Of note, for 11 of the 12 network patches, within-individual similarity was found to be greater than between-individual similarity (Figure 3) – suggesting that despite similarities, consistent inter-individual differences exists.The salience patch was the notable exception. This patch could not be successfully assigned, largely due to the small size of the salience network and its spatial variation in individuals.

### Individual Areal Boundaries Have Familial and Genetic Associations

To demonstrate the feasibility of relating measures of similarity for areal organizations to known sources of inter-individual or phenotypic or biological variation, we made use of the R-fMRI dataset (Yang et al., 2016) generated as part of QTIM (de Zubicaray et al. 2008). Specifically, we examined whether inter-individual differences in full-brain areal measures (gradient maps, edge maps) could be related to familial (i.e., sibling vs. non-sibling pair) or genetic (monozygotic twins vs. dizygotic twins) relationships. To accomplish this, for every pairing of individuals in the sample, we calculated the spatial correlation between the iFC similarity-based gradients maps, as well as for the edge maps. Our a priori predictions were that gradient/edge maps would be more similar among siblings than nonsiblings, and more similar among monozygotic twin pairs than dizygotic twin pairs. As expected, the correlation between siblings was significantly higher than non-siblings (gradient: p<10^−23^; edge map: p<10^−50^) (See Figure 4). The correlation between MZ pairs was significantly higher than DZ pairs in edge (p=0.002), and marginal significant in gradient (p=0.075).

**Figure 4.**
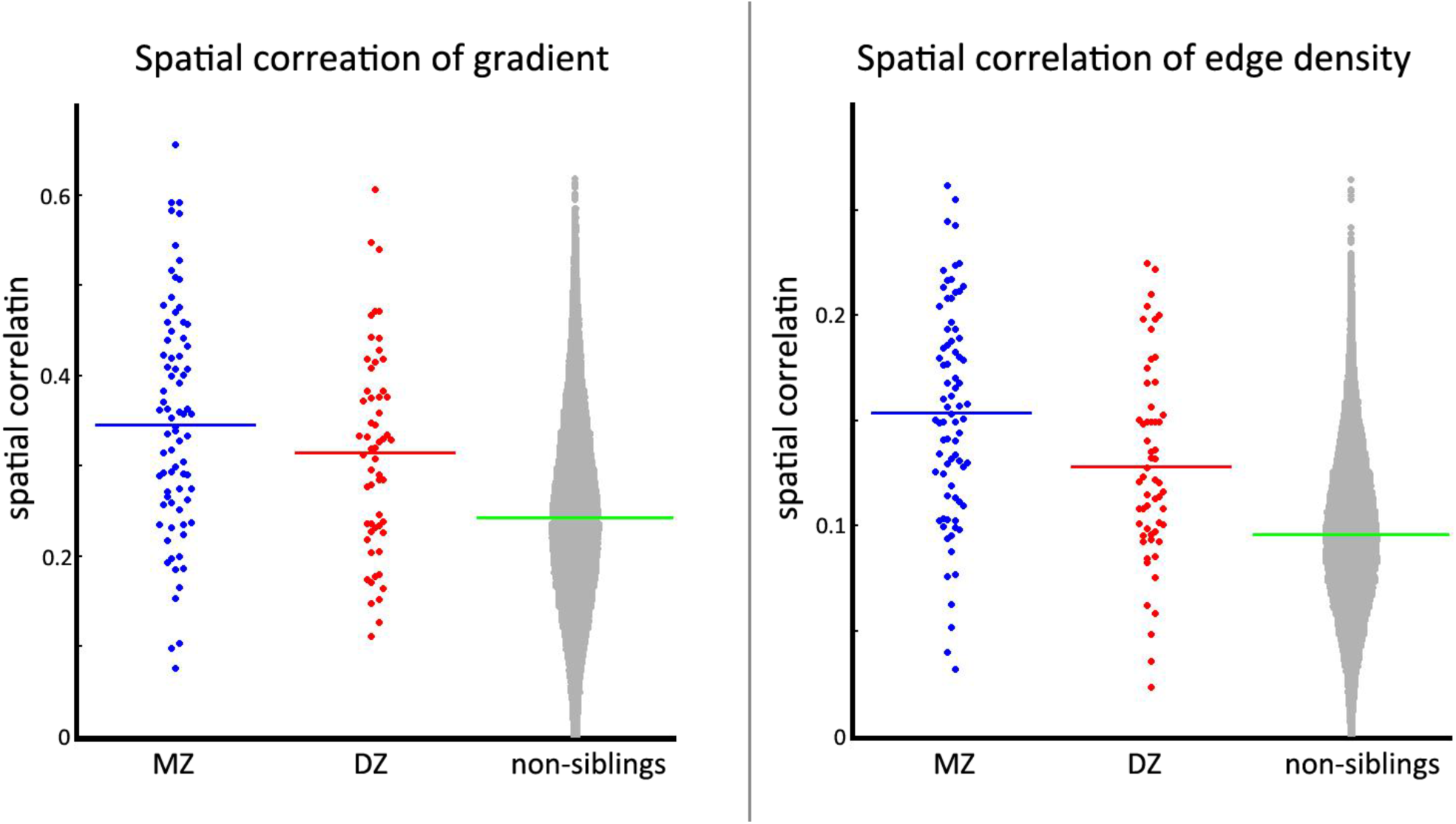
Spatial correlations of gradients and edge density in iFC similarity between MZ, DZ, and unrelated pairs. Blue lines are the mean correlation across MZ pairs for gradient (A) and edge density (B). Red lines are the mean correlation across DZ pairs for gradient (A) and edge density (B). Green lines are the mean correlation across all unrelated pairs of QTIM dataset for gradient (A) and edge density (B).

Given that a trend was found between MZ and DZ pairs with respect to the level of spatial similarity for anatomical images (p=0.08), we further tested whether the observed heritable functional transitional profiles could have been confounded by structural information. However, we did not find any significant correlations between similarity of structure and gradient/edges in twin pairs (all p>0.3). The significance comparisons between MZ and DZ pairs showed almost identical p-values after being controlled the structural similarity (gradient: p=0.072, edge: p=0.002).

### Data Requirements for Mapping Inter-Individual Differences in Areal Organization

Recent work has emphasized the value of using longer data acquisitions (or combinations of shorter ones) to increase the stability of estimates of functional connectivity. In particular, using a correlation matrix based on 380 minutes of data from a single participant as a reference, Laumann et al. 2015 found that estimates of the full-brain correlation matrix did not plateau in consistency (with respect to the reference) until 27 minutes of data were included. In the present work, we attempted to further inform this issue by examining the stability of gradient and edge maps, as data is added in 10-minute increments. More specifically, first, for each participant, we randomly selected 5 of the 10 sessions and used their data to derive reference gradient and edge maps. Then, for each participant, we randomly selected samples of 10, 20, 30, and 40 minutes of data from the remaining five 10-minute sessions, for spatial correlation of the resultant gradient/edge maps with the same participant’s 50-minute reference maps.

Regardless of whether 10, 20, 30, 40 or 50 minutes of data were used, the resultant gradient or edge map for an individual consistently showed a higher spatial correlation with the reference map calculated for the same participant, than with the reference map for any other participant. In other words, for all participants, one could readily distinguish whether the map derived was properly matched with that from the same participant as opposed to different participants. The distribution of the spatial correlations to the reference gradient or edge map is demonstrated in Figure 5.

**Figure 5.**
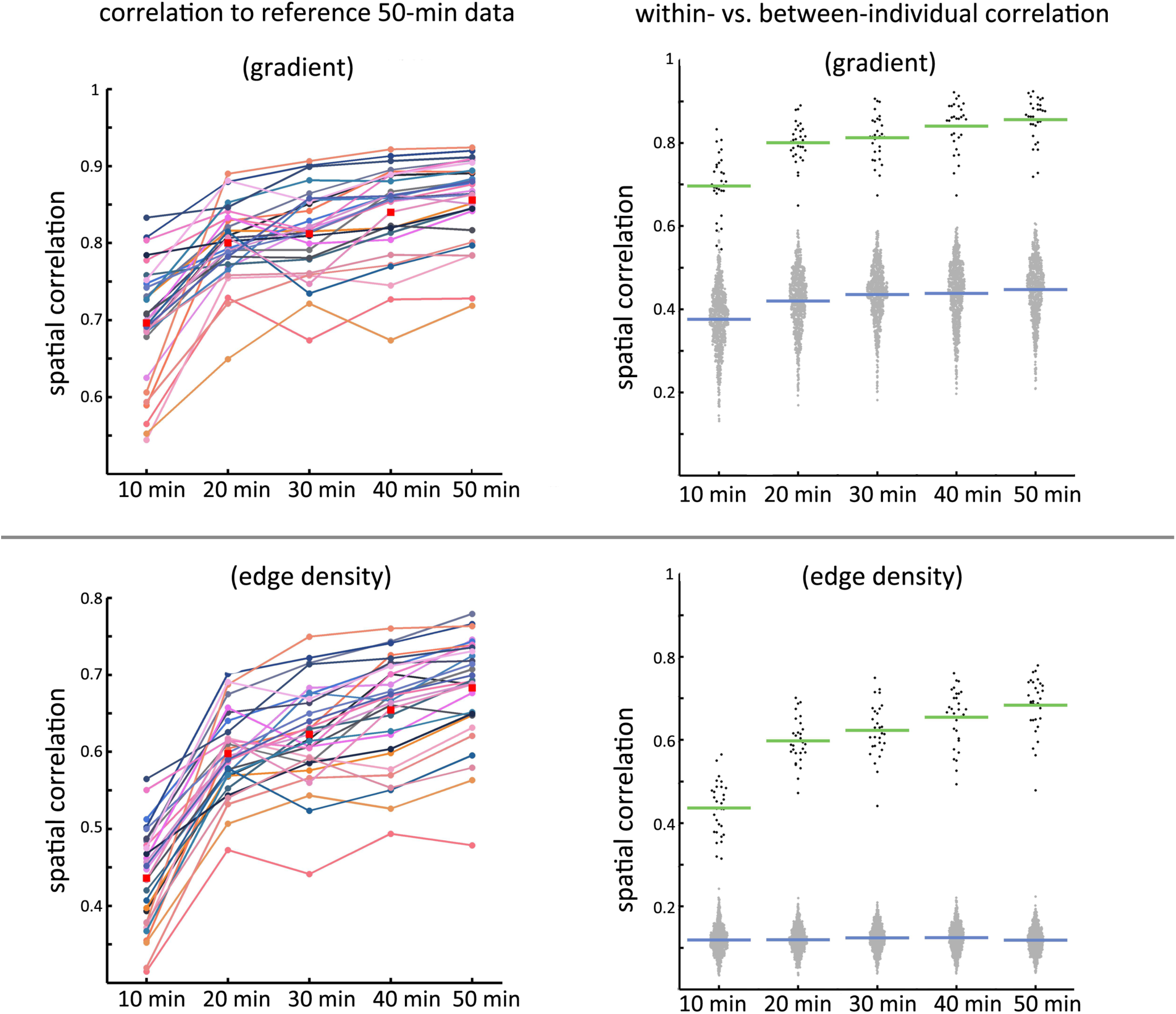
The stability of individual gradient and edge maps as a function of the amount of data used for gradient estimation. A: Each dotted-line represents a different participant; the spatial correlation between maps generated from each incremental amount of data (10, 20, 30, 40 and 50 minutes) and those from the remaining 50 minutes session data (reference) are depicted (top: gradient; bottom: edge density). Red dots are the mean correlation across participants at each data subsets B: The distribution of within-individual correlations (black dots) and between-individual correlations (grey dots) at selected data subsets to the 50-minutes references for each individuals.

It is important to note that the consistency of findings obtained across the various quantities of data should not be taken to infer that there were no differences. Consistent with predictions based upon prior work, across participants, each 10-minute increment in data produced an incremental increase in the mean correlation with the reference images – for both gradient and edge maps (Figure 5A). Focusing first on gradient maps, we found that the average within-participant correlation between the map generated from a single scan session (10 min, 295 time points) and an individual’s reference image was r=0.70 (SD=0.08); this progressively increased to a high of r=0.86 (SD=0.05) when data from five sessions (50 min, 1475 time points) was used (Figure 5B, upper). Statistical testing found that the observed increase was significant for each 10-minute increment (p<0.001), except for the 20- to 30- minute increment (p=0.108). For edge density, we found that the average within-participant correlation between the map generated from a single scan session and an individuals’ reference image was r=0.44 (SD=0.06); this progressively increased to a high of r=0.68 (SD=0.07) with 50 min (1475 time points) (Figure 5B, lower). Statistical testing found that increases in correlation for each 10-minute increment were significant (all p<0.001).

To gain greater insights into the influence of scan time in terms of test-retest reliability, we calculated the ICC of gradient and edge density maps for each of the scan durations of 10 min, 20 min, 30 min, 40 min and 50 min by randomly selected 1, 2, 3, 4, or 5 10-minute sessions of the 10 available sessions for each participant (i.e., 10, 20, 30, 40 or 50 minutes of data, respectively). As shown in Figure 6, as a measure of functional local transition, the gradient of iFC similarity can achieve good test-retest reliability (ICC > 0.5 for over 50% vertices across cortex) with a scan duration of 20 min, which is consistent with prior work (Laumann et al., 2015). A relationship was observed between gradient strength and ICC across vertices (see Supplementary Figure S5).

**Figure 6.**
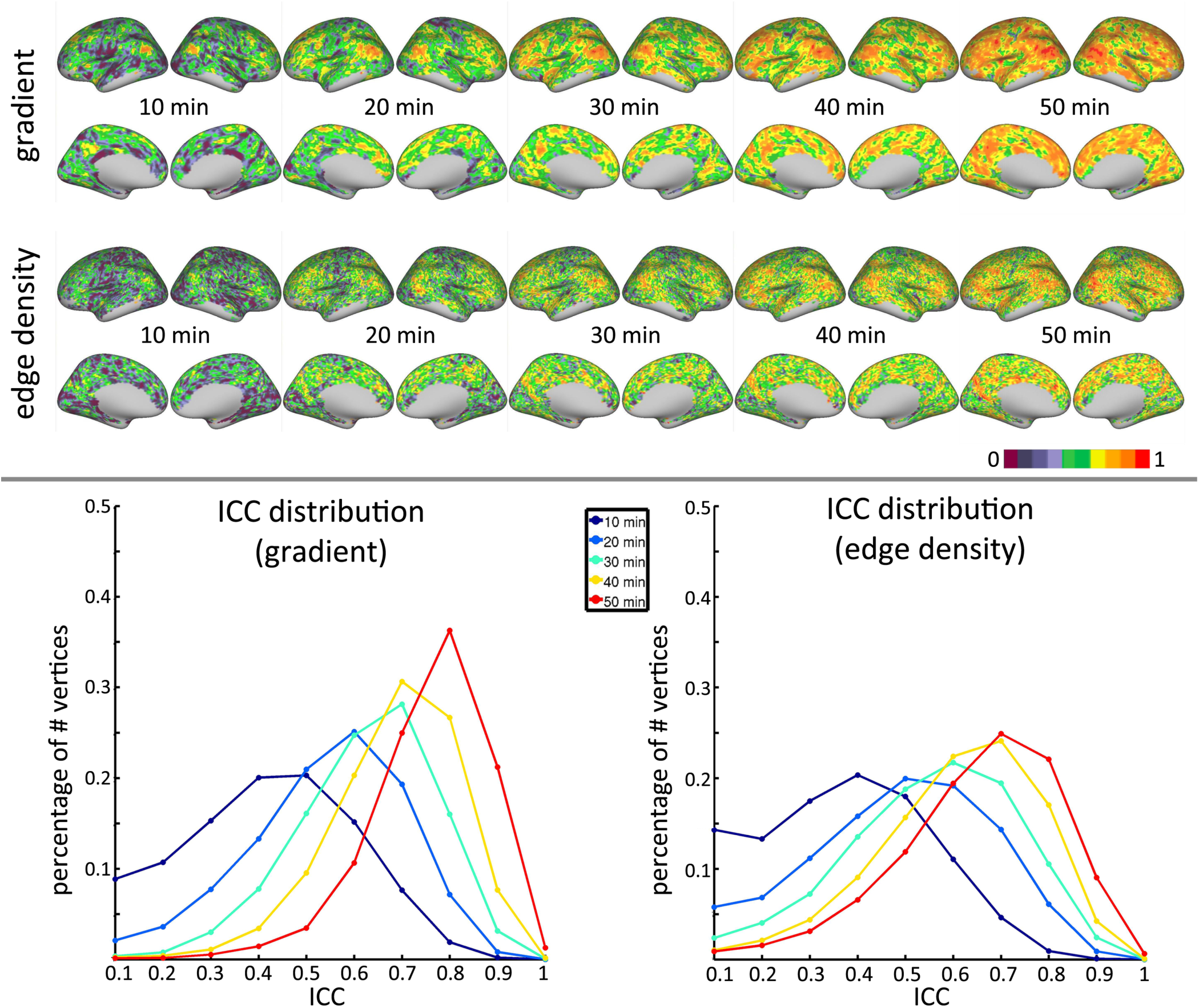
The influences of scan time on test-retest reliability of gradient and edge density of iFC similarity. The reliability of gradient and edge density were estimated as intra-class correlation maps for each of the scan durations of 10 min, 20 min, 30 min, 40 min and 50 min (top panel). To access the influences of scan durations on the ICC distribution, the histograms of ICC scores are plotted for each scan durations in the bottom.

To provide a single measure of reliability, we complemented the spatial correlation-based measures of repeatability with image intra-class correlation coefficient (I2C2) (Shou et al. 2013). The I2C2 was increased with scan time for gradient (10 min: 0.42, 20-min: 0.54, 30-min: 0.61, 40-min: 0.67, 50-min: 0.73) and edge density (10 min: 0.35, 20-min: 0.48, 30-min: 0.55, 40-min: 0.61, 50-min: 0.65). In addition, we replicated the I2C2 findings in eNKI test-retest dataset (multiband EPI sequence TR=645ms, 10 min, 895 time points). The I2C2 was nearly identical to that observed for the 10-minute HNU dataset (0.49 for gradient and 0.34 for edge density). However, the distributions of vertex-wise ICC values for gradient and edge maps derived from the eNKI-TRT were superior to those observed for the 10-minute HNU scans (gradient maps: mean ICC [eNKI-TRT] = 0.40 [SD=0.22], mean ICC [HNU] = 0.37 [SD=0.18]; edge density: mean ICC [eNKI-TRT] = 0.30 [SD=0.21], mean ICC [HNU] = 0.32 [SD=0.18]); see supplementary Figure S6.

### Within- and Between-Individual Variability

It is generally accepted that anatomical components of higher order association areas tend to have greater anatomical variability among individuals than those comprising lower-order sensory and motor areas (Mueller et al. 2013). To gain similar insights into potential regional variation in areal organization, we examined the within- and between-participant variation separately. Figure 7A and 7B depict the percentage of the total variability that is the intra- vs. inter-individual variability (after accounting for that attributable to nuisance signals) for each, gradient maps and edge maps. Intra-individual variability indexes the temporal stability of transition scores within participant while inter-individual variability indexes the stability of transition scores across the participants. Both intra- and inter-individual variability demonstrated a non-uniform distribution across brain regions. For both gradients and edges, between-individual differences were largest in high-order association cortex including the lateral prefrontal lobe, lateral parietal lobe, and around the border of the PCC-precuneus default mode networks, while lower in sensory-motor cortices and medial occipital visual cortices. Within-individual variability depicted the complementary pattern from interindividual variance, with the maximal variation of gradient and edges being noted within sensory-motor and visual regions. Regions within the defaultmode network demonstrated a moderate level of within-individual variability.

**Figure 7.**
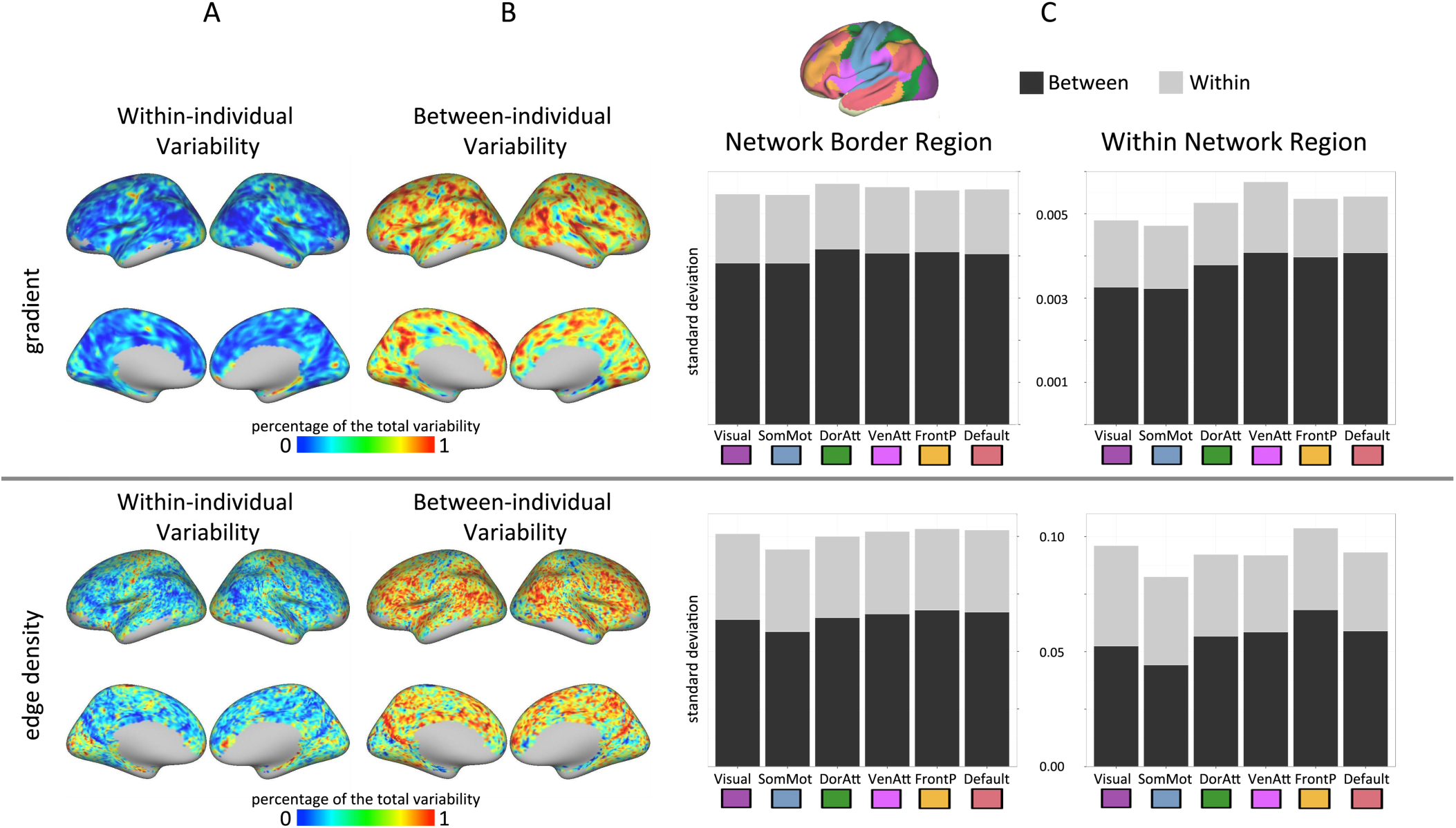
Within- and between-individual variability for gradient and edge density maps based on iFC similarity. A: The percentage of within individual variation (relative to total variation) in gradients (top) and edge density (down) scores (based on IFC similarity) across sessions; B: The percentage of between individual variation in gradient (top) and edge density (down) scores (based on iFC similarity); C: Within- and between- individual variation (standard deviation) at network borders (defined as confidence <0.3 along the network borders) and within network region (defined as confidence >= 0.3 within each network). The limbic network was not included here due to the substantial signal loss in temporal areas.

To provide a network-level perspective of intra-individual variations in gradient and edge densities we made use of the confidence maps for the 7 networks in Yeo et al. (2011)^1^. Specifically, the standard variations were averaged at network borders (defined as confidence <0.3 along the network borders) and within network region (defined as confidence >= 0.3 within each network). Without exceptions, the between-individual differences contributed more than 60% of the variance in transitions at the border region of 7 networks (Figure 7C). For the within network regions, the between-individual differences still accounted for more variance than within-individual differences, however, the somatomotor and visual networks exhibited an increases of within-individual variation to 45% and 46%, respectively (Figure 7C). This suggested the spatial iFC profile in the somatomotor and visual networks were more state dependent than the other networks. This finding is consistent with that of Mennes et al. (2010), Craddock et al. (2013), and Laumann et al. (2015).

### Transition Pattern Based on Local-, Global, and Network-Scale iFC

Previous studies using stimulus-based functional neuroimaging methods have suggested that areal discrimination maps could vary depending on the specific property or properties used in their definition, e.g. angular and eccentricity representation maps for distinct areas of early visual cortex (Buckner and Yeo 2014)(Buckner and Yeo 2014)(Buckner and Yeo 2014)(Buckner and Yeo 2014)(Buckner and Yeo 2014)(Buckner and Yeo 2014)(Buckner and Yeo 2014)(Buckner and Yeo 2014)(Buckner and Yeo 2014)(Buckner and Yeo 2014)(Buckner and Yeo 2014)(Wandell and Winawer 2011; Buckner and Yeo 2014). Here, we examined whether transition patterns are dependent upon the specific intrinsic brain measure used for their definition. Specifically, we repeated our analyses focused on the definition of gradient and edge maps using commonly examined measures in the literature (i.e., ReHo, DC, EC, and DR-Networks). We then measured the convergence of transition patterns for both gradients and edge density from iFC similarity, ReHo, DC, EC and DR-Networks. Figure 8 demonstrates the average pairwise correlations of gradient between all functional metrics. Among all the metrics, the centrality maps (DC and EC) showed the most remarkably similar transition scores (r=0.94, SD=0.03). Edge density of iFC similarity exhibited low correlation with ReHo (mean r=0.11, SD=0.08) but had notable correlations with DC (mean r=0.30, SD=0.19), EC (mean r=0.45, SD=0.18), DR-medVis (mean r=0.26, SD=0.15), DR-occVis (mean r=0.25, SD=0.13), DR-LatVis (mean r=0.43, SD=0.14), DR-DMN (mean r=0.60, SD=0.11), DR-Cerebral (mean r=0.32, SD=0.13), DR-SenMot (mean r=0.64, SD=0.10), DR-Auditory (r=0.61, SD=0.12), DR-Control (r=0.47, SD=0.12), Dr-FrontL (r=0.49, SD=0.08), and DR-FrontR (mean r=0.51, SD=0.10). These findings are more reflective of global- and network-level iFC characteristics, than local features (Figure 8). The Kendall Coefficient of edge density for all the functional indices was 0.29, SD=0.05. Similar finding were observed for edge density (Figure S7).

**Figure 8.**
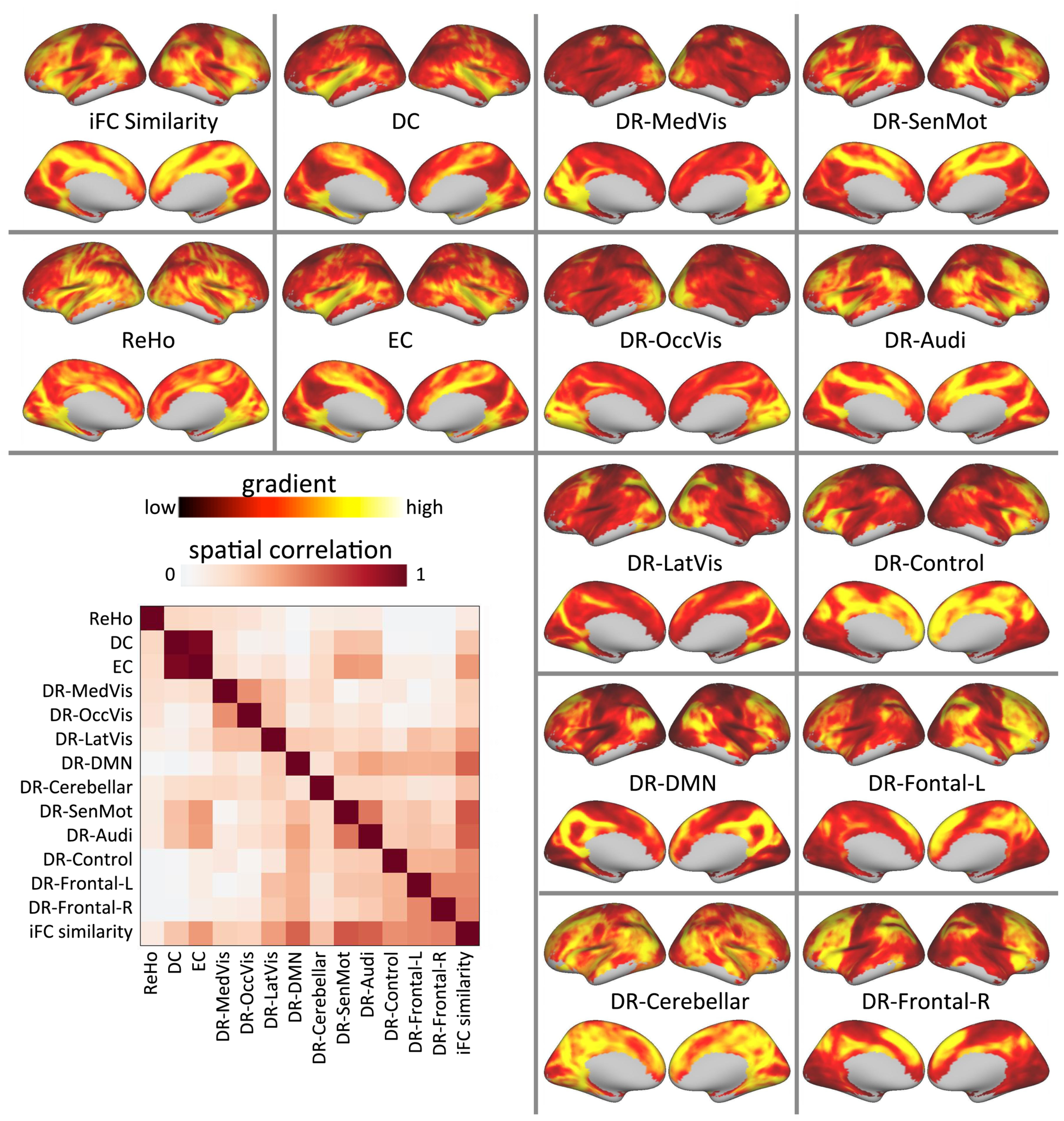
Group-level gradient maps for each functional indices from two 50-minutes subsets. For each participant, we calculated the spatial correlation matrix of the gradient maps for various functional indices, and then averaged across individuals to provide the spatial correlation matrix (bottom left corner). The group-level edge density maps were demonstrated in Supplementary material in Figure S7.

### Within- Versus Between-individual Variance of Transition Pattern based on Local-, Global, and Network-Scale iFC

Reminiscent of the finding for iFC-similarity above, for all the functional metrics, gradient (and edge) maps calculated from different sessions in the same individual exhibited a consistently higher amount of spatial correlation with one another than those from differing individuals (Figure S8). However, with the exceptions of ReHo, DR-DMN, and DR-FrontoL, the spatial correlation of gradient and edge maps generated in different sessions for an individual were generally lower than iFC similarity. Those measures characterized by a higher between-participant correlation in gradient or edge maps, typically exhibited a higher within-individual correlation (i.e., repeatability) over time. Given that the gradient of iFC similarity was averaged from 20k gradient maps across vertices, this may in part reflect the cleaning of data within an individual by averaging.

### Reliability of Transition Properties based on Local-, Global, and Network-Scale iFC

While our analyses primarily focused on full-brain transition patterns, we did take the opportunity to provide insights into the test-retest reliability of vertex-based gradient and edge map scores. Specifically, we calculated vertex-wise intra-class correlation coefficients for each, gradients and edges, using the two 50-min subsets. Gradients in iFC similarity showed considerably high reliability across cortex, especially in PCC-DMN, LP-DMN, and frontoparietal association cortex (Figure 9). The pattern was more apparent in reliability of edge maps (Figure S9). Intriguingly, highly reliable edges were found around the border of PCC-DMN, LP-DMN, and the frontoparietal regions, which were observed as a common transition zones in variant iFC metrics. Among different functional indices, ReHo exhibited the highest ICC in transition scores. The intra- and inter-individual variation analysis confirmed that high ICC of ReHo was due to the considerably small amount of intra-individual variation. For DR networks, the ICC of gradient and edge scores showed non-uniform spatial distribution in different networks. The vertex-wise ICC of gradient/edges scores obtained for a given DR network tended exceptionally high when looking at vertices that were part of the network (Figure 9). Similarly, in contrast to iFC similarity, vertex-wise ICC was strongly related to gradient strength for the nearly all of the DR networks (the DR Cerebellar network is an exception) (see Supplementary Figure S10).

**Figure 9.**
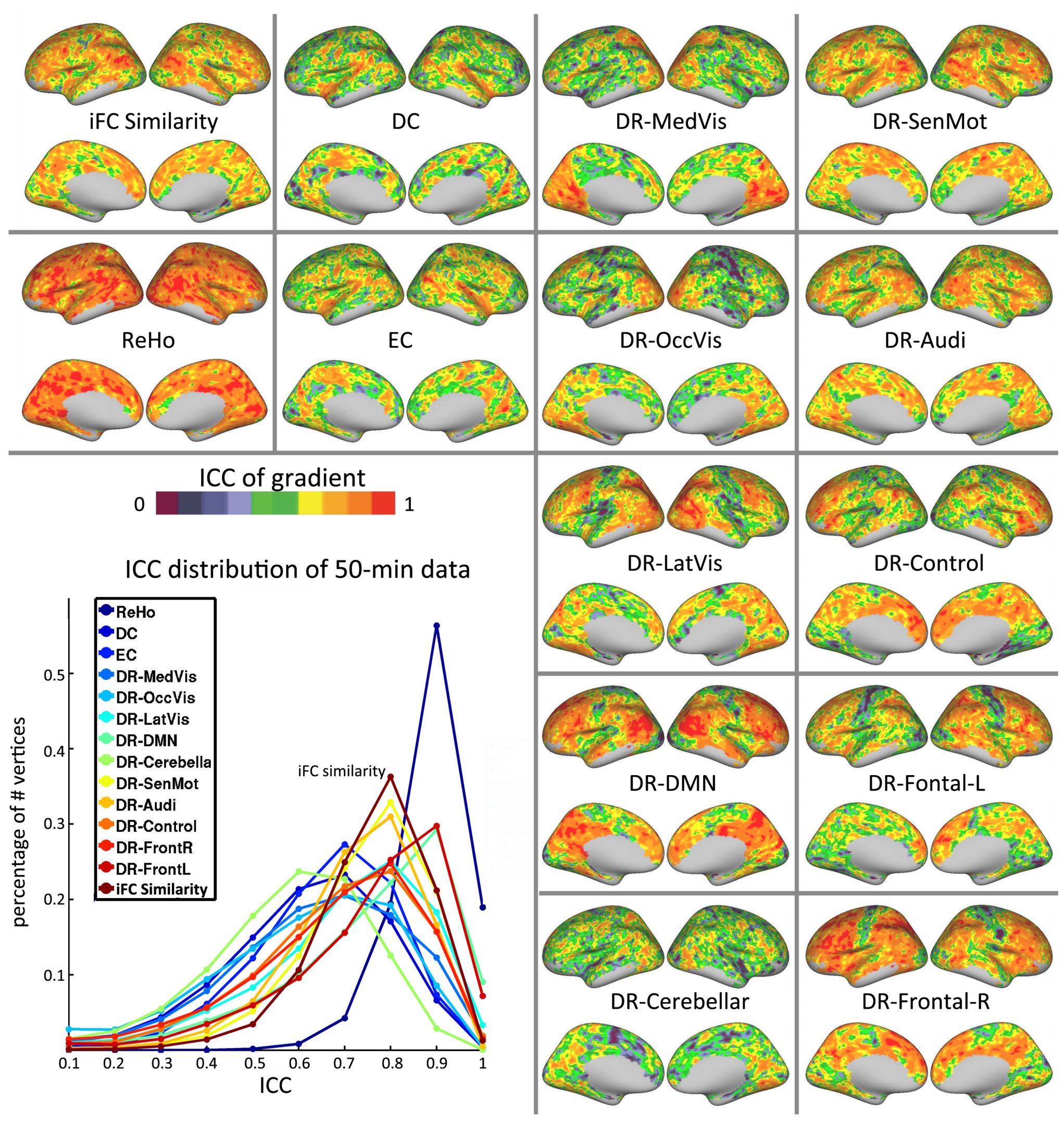
The intra-class correlation maps for the gradient of 14 functional indices are depicted here (ICC was calculated between two 50-minute subsets of data generated through random selection [without replacement]). The distribution of ICC for edge density maps for each functional metric is shown in the bottom-left corner. The ICC maps for edge density maps were demonstrated in Supplementary material in Figure S9.

### Parcel Homogeneity based on Local-, Global, and Network-Scale iFC

In evaluating and comparing the optimality of parcellation strategies, unit homogeneity is commonly viewed as a desirable property. As such, we opted to calculate the homogeneity of full-brain iFC patterns among voxels within a given parcel as a means of comparing the parcellations produced using the differing R-fMRI measures. More specifically, we first computed the grey matter (cortical and subcortical) connectivity map at each vertex using the reference 50-min data. Then for each parcel within a given brain parcellation from the other 50-min data, the Kendall’s coefficient of concordance was calculated from the functional pattern of all vertices within a parcel. We then averaged the KCC-homogeneity scores for each approach at the individual-level and test which approach performed most accurate parcellation. Figure 10 demonstrated the KCC-homogeneity scores across all the participants. Locally Weighted Scatter-plot Smoother (LOWSS) was applied to demonstrate the average KCC-homogeneity for each of the functional indices. Parcellations generated using the iFC similarity methodology achieved the highest within-unit homogeneity, followed by the DR-network. Parcellations generated using ReHo and DC exhibited the lowest homogeneity. This suggested that local- and network-scale measures might not be well suited at generating homogenous parcels for depicting the complexity of cortical areal organization as a whole system.

**Figure 10.**
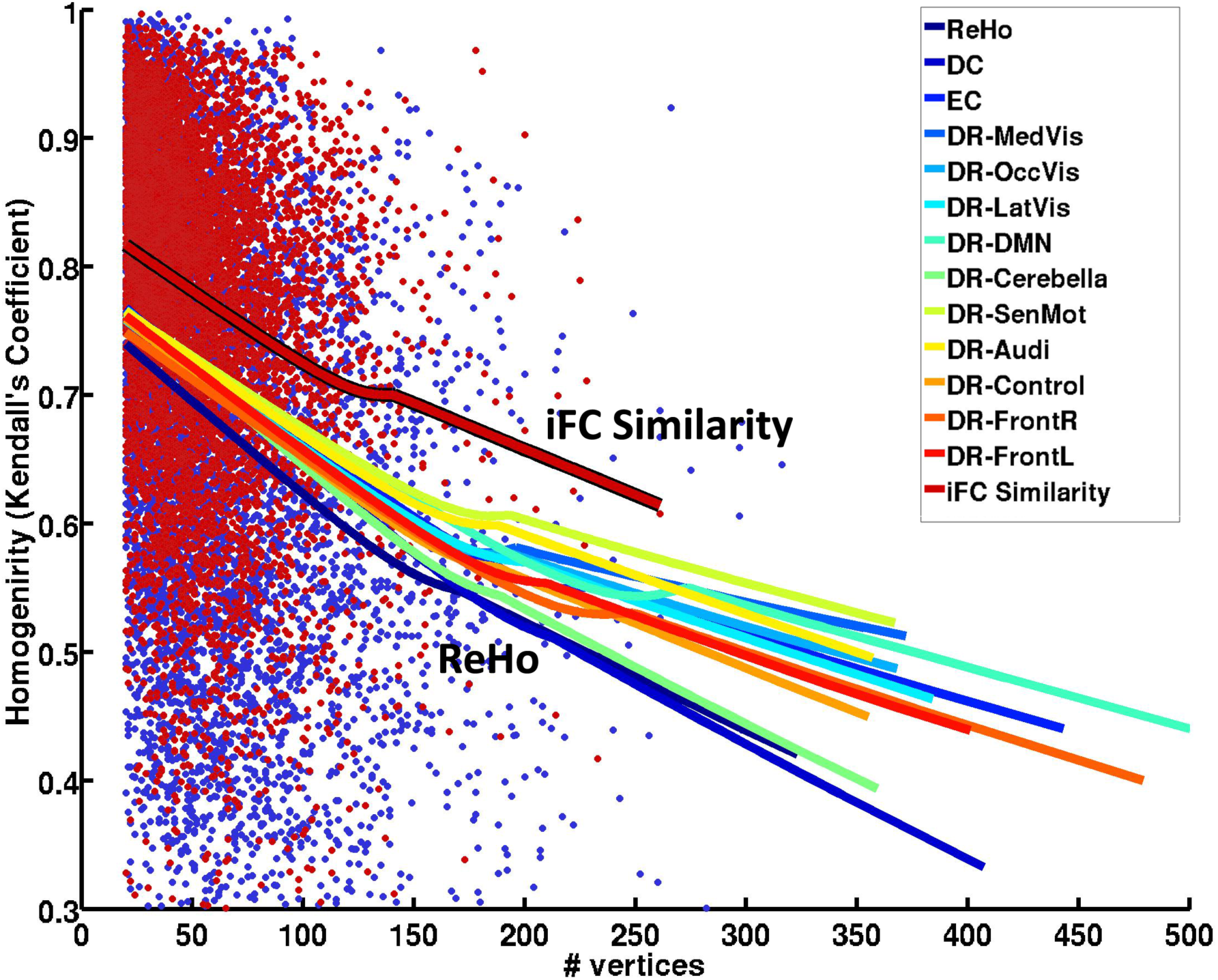
Homogeneity of individual parcels by parcel size based on 14 iFC indices. Red dots are homogeneity scores for each parcel driven from iFC similarity across individuals. Dark blue dots are homogeneity scores for each parcel based on ReHo across individuals. Each line is the LOWESS fit represents the effect of parcel size on homogeneity across individuals.

### Effect of Surface Geometry and Surface Registration

A potential concern regarding the methodologies presented is that the underlying surface geometry may influence the presence and locations of estimated gradients and edges. To test for this, we repeated our gradient-based analyses using surrogate fMRI data that consisted exclusively of white Gaussian noise. Figure 11A–B summarizes the spatial correlation of iFC similarity and its gradient between real and random data (ordered by participants and scan sessions, real to random). For the surrogate data, within-participant spatial correlations were again higher than between participant spatial correlations. However, spatial correlations of gradients between maps derived from real and surrogate R-fMRI showed a relatively low degree of correlation – even for the same participant (gradient: mean r=0.03, SD = 0.06). These findings suggest that the surface geometry explained only less than 1% of variance in the gradient of iFC similarity.

**Figure 11.**
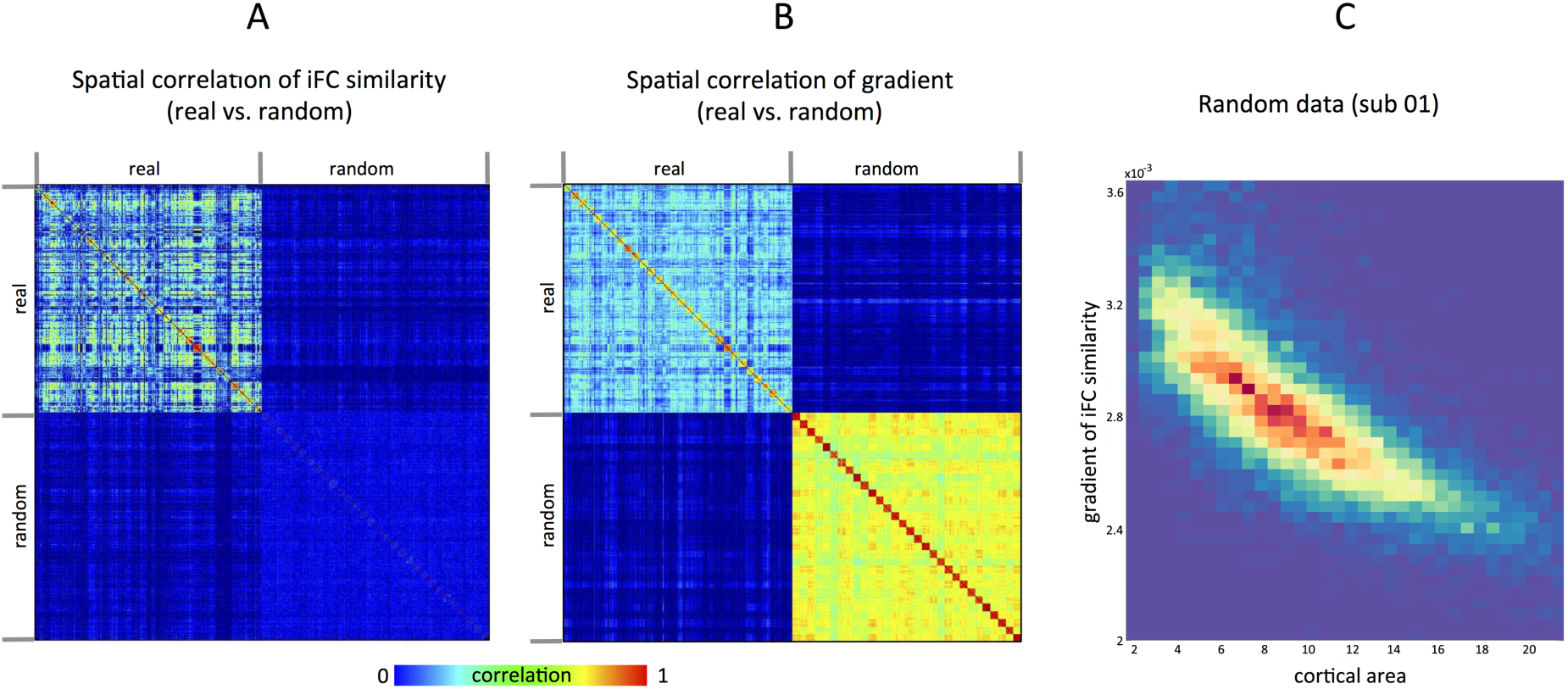
Effect of surface geometry and surface registration. A: The spatial correlations matrix of mean iFC similarity from real and random data. B: The spatial correlations matrix of mean gradient in iFC similarity from real and random data. The correlation matrixes are ordered by participant: top left sub-matrix is from real data (first ten rows for ten sessions of participant 1, second ten rows for ten sessions of participant 2, and so on) and bottom right sub-matrix is from random data in the same order. C: The 2D histogram depicting the relationship between cortical area and mean gradients of iFC similarity computed from random data.

Thus, while inter-individual variations in the surface geometry in and of themselves can produce repeatable gradients, they are unrelated to findings observed with functional data. Although beyond the scope of the present work, preliminary analyses suggest that the strength of the findings obtained for surrogate data is largely attributable to inter-individual variations in the distance between adjacent neighbors, which impacts gradient computation (Figure 11C). However, it is worth noting that the gradient values from noise were generally minute; for nearly all vertices, the real gradient score exceeded the 90th percentile of scores generated from noise. In those cases where this was not the case, the failure to exceed the 90th percentile was due to the presence of a relatively low gradient score in the real data (thus failing to rise above the noise) (see the supplementary Figure S11).

To address any concerns about the possible effects of noise with regard to comparison of within- and between-individual variance, we repeated our analyses focused on establishing the similarity of gradient maps across subjects and sessions; though this time, we regressed out noise-based gradient maps for each before testing the differences of within- versus between-individual variance. The correct identification rate was almost the same as 90.44% for real data and 90.96% after regressed out the noise-based gradient. In sum, the whole transition patterns were predominantly driven brain function, rather than structure.

As a final sanity check, we repeated our edge detection analyses using the similarity of iFC maps rather than gradient maps. Spatial correlations between sessions (for the same participant) were quite low – again, reinforcing the validity of the findings obtained in our primary analyses.

## DISCUSSION

Using publically available datasets, the present work assessed the ability of gradient-based iFC boundary mapping approaches (Cohen et al. 2008; Wig et al. 2013, 2014; Gordon et al. 2014, 2015; Laumann et al. 2015) to reliably characterize inter-individual variation in the transitional properties of cortical areas throughout the brain. First, from the perspective of “finger-printing”, across the 10 scan sessions, we were able to demonstrate the ability to accurately identify R-fMRI data obtained from a given individual based upon the similarity of full brain gradient and full-brain edge maps derived from each of the scan sessions. Supplementary analyses demonstrated that these abilities were not driven by structural contributions. Second, using image intraclass correlation coefficient (I2C2), we were able to demonstrate moderate full-brain test-retest reliability for a 10-minute scan; I2C2 progressively increased with the amount of data available, reaching relatively high levels when 50 minutes of data were used. Third, using vertex-wise intraclass correlation coefficient, we demonstrated low to high testretest reliability depending on region; again, as the amount of data included was increased, ICC values approached impressively high levels. Of note, within-participant variation in transition zone properties appeared to be greater in lower order networks than higher order, while between-individual variation was greater in higher order networks; these findings are consistent with those of prior efforts examining iFC profiles. Intriguingly, using a previously published twin dataset, the present work was able to push this finding one step further, showing associations between the similarity of edge maps and each, familial relations (i.e., sibling vs. non-sibling) and genetic (i.e., monozygotic vs. dizygotic twin pairs). Finally, the present work extended prior work by employing alternative iFC measures to define gradients, revealing similar but distinct gradient profiles for each of the iFC features – all of which exhibited impressive repeatability and reliability over time if sufficient data is used. Importantly, the various findings obtained in the present work did not appear to be related to potential sources of artifactual variation, such as head-motion or surface registration. As such, our findings increase confidence in the potential for gradient measures to be used as a novel feature upon which interindividual differences in brain function can be mapped and eventually related to phenotypic variation.

### Individual Areal Organizations Are Unique

Long focused on the comparison of groupings of individuals that differ on one or more features (e.g., diagnostic status), the delineation of inter-individual variation has emerged as a central focus in emerging neuroscientific and clinical agendas (e.g., biomarker identification). Several studies have emphasized the potential utility of taking variations in cortical area transition zone properties into account when attempting to catalog phenotypic variation (e.g., Di Martino et al. 2009; Adelstein et al. 2011). Our analyses revealed full-brain transition patterns on an individual basis, which distinguished participants from one another across scan sessions. Findings that the topological architecture differs between participants are consistent with recent work highlighting inter-individual variation in key features of the topological architecture (e.g. area size and shape) (Gordon et al. 2015). Perhaps more interesting, were findings that full-brain gradient and edge maps can distinguish nearly all individuals using only 10 minutes of data (Figure 1C). This finding echoes recent studies, which suggested the potential for full-brain characterizations of the intrinsic brain to ‘fingerprint’ individuals (Poldrack et al. 2013; Miranda-Dominguez et al. 2014; Finn et al. 2015) – a capability that is encouraging for efforts towards biomarker discovery. Although not a focus of the present work, confirmatory analyses replacing actual functional data with random noise were able to demonstrate the fingerprinting abilities of structural brain properties embedded in the cortical surface as well. Importantly, these structural fingerprints were largely unrelated to the various indices derived from true functional data. Taken together with prior work, it appears that future work may find potential value in the development of fingerprinting profiles.

### Individual Functional Areal Organizations Are Reliable With Sufficient Scan Time

As suggested by prior work, more is generally better when discussing the test-retest reliability for RFMRI findings. While fingerprinting could be performed with relatively high accuracy using only 10 minutes of data, the bar for drawing reliable functional boundaries appears to be higher. Recent work using a highly sampled individual dataset found that 27-min was required for connectivity scores to plateau in large-scale iFC networks (Laumann et al., 2015). Consistent with this report, we found that 20 minutes or more of data were required for gradient and edge density scores to achieve moderate to high test-retest reliabilities at the majority of vertices.

A key caution is that while studies attempting to determine the minimum sufficient data required to optimize an analysis commonly report their findings in terms of time, most do not allow this construct to be disentangled from the number of time points included. In our findings, although the number of time points for eNKI-TRT dataset (TR=645 ms, 10 min, multiband sequence) was about three times to HNU dataset (TR=2000 ms, 10 min, standard sequence), only the vertex-wise reliabilities showed an advantage over the 10-minute HNU dataset, and they were inferior to those obtained with 20 minutes of HNU data. This may suggest the amount of time sampled could be more important than the number of timepoints. However, it is important to note that the eNKI-TRT dataset used an early version of the CMRR multiband sequence, rather than standard EPI as in the HNU dataset. Future work will be required to disambiguate the various factors further. It is also possible that continued refinement of preprocessing strategies may help to further decrease data needs for such analyses. Nonetheless, we emphasize that at the present time, it appears prudent to obtain more data rather than less if the goal of a study is to reliably characterize fine-grained functional boundaries.

### Functional Boundaries Have Familial and Genetic Associations

Recent work has begun to explore genetic and environmental influences on inter-individual variation in indices of intrinsic brain function (Glahn et al. 2010; Schutte et al. 2013; Fu et al. 2015; Yang et al. 2016). Already, connectivity within a number of functional has been shown to have moderate to high heritability (e.g. default, fronto-parietal, somatosensory, visual, and attention networks). Complementing these findings are those of studies suggesting the heritability of some between-network functional connectivity (e.g., Yang et al., 2016), as well as larger global network architecture properties (e.g. modularity, clustering coefficient, global efficiency) (van den Heuvel et al. 2013; Sinclair et al. 2015). The present work helps to build upon this growing body of literature by suggesting possible familial (sibling vs. non-sibling) and genetic (monozygotic vs. dizygotic) associations for measures of areal brain organization. Importantly, these associations did not appear to be driven by the underlying neuroanatomy. Consistent with previous studies suggesting that different genetic mechanisms are responsible for the structure and function of brain areal organization (e.g., Glahn et al. 2010). Of note, familial effects (sibling vs. non-sibling) were found to be relatively robust, suggesting potential environmental contributions as well – a finding consistent with prior work (e.g., Yang et al. 2016)

In light of the relatively limited amount of R-fMRI data available per participant in this QTIM dataset (scan duration = 5 minutes), the results obtained were taken to be particularly encouraging. We considered pursuing quantification of heritability for the indices of areal organization examined in the present work, we decided against it as: 1) comprehensive exploration of the topic was beyond the scope of the present work, and 2) the limited data available for each participant is still likely to compromise the quality of gradients and edge maps obtained, thereby leading to an underestimation. Future work with larger datasets focused on the collection of large amounts of R-fMRI data from each participant (e.g., the Human Connectome Project, Van Essen et al. 2013) would be more appropriate for such determinations.

### Sources of Within- and Between-individual Variability in Areal Organization

A growing number of studies are appreciating regional and network-level differences in the stability of connectivity patterns across individuals and time, which can be informative to our understanding of brain development and function. Intra-individual variability characterizes the temporal stability of transitional zone properties for cortical areas (i.e., low variation infers high stability), while interindividual variability captures the conservation of these zones from one person to the next (i.e., low variation infers conservation). Intra- and inter-individual variability of transitions in iFC features were not uniformly distributed across the cortex in the present work. Specifically, multimodal association networks (e.g., default, dorsal attention, and executive control) exhibited greater variability between individuals than within; in contrast, unimodal networks (e.g., visual, sensorimotor) were found to have lower inter-individual variability whilst high intra-individual variability. These results echoed previous findings highlighting greater intra-individual variation in lower-order networks (e.g. Mennes et al. 2010; Craddock et al. 2013), whose functional interactions are more heavily influenced by current task demands (Mennes et al. 2010). Similarly, they echo the findings of studies suggesting greater interindividual variation in higher order networks (Mueller et al. 2013; Wang and Liu 2014; Langs et al. 2015; Laumann et al. 2015), which appear to be more affected by genetic and environmental factors (Anderson and Finlay 2014; Gao et al. 2014). The greater susceptibility of higher order multimodal association networks to environmental influences is not surprising given their more protracted developmental period relative to unimodal (Mueller et al. 2013; Zilles and Amunts 2013). Additionally, the later evolutionary development and enlargement of association cortices supporting higher-order network function may contribute in part to our findings of increased variation (i.e., decreased conservation) across individuals (Van Essen and Dierker 2007; Brun et al. 2009; Van Essen, Glasser, Dierker, and Harwell 2012; Chan et al. 2014).

Potential confounds that can arise in consideration of inter-individual variation in functional transition zones come from differences in cortical folding patterns (Hill et al. 2010). For example, sulcal depth can exhibit patterns of inter-individual variation that are similar to those observed in iFC (Mueller et al. 2013). Additionally, given that current surface-registration algorithms are based on the anatomical curvature, the cross-individual registration could lead to non-uniform misalignment across cortex, thereby contributing to individual variability into functional transitions (Robinson et al. 2014). In the present study, we explored these potential confounds through the replacement of functional MRI data with random noise. The underlying anatomical architecture had only a slight relationship on functional transition profile of iFC features (Figure 8B), reaffirming confidence that our findings are driven by iFC.

### Different Functional Features Shows Similar but Distinct Areal Transition Profiles

Recent years have witnessed the emergence of a growing number of R-fMRI measures, each capturing a unique aspect of the intrinsic functional architecture. The distinctions among some indices can be readily delineated based upon differences in their definitions (e.g., centrality measures) or the networks being examined (e.g., dual regression of network components); while for others, the exact positioning of one measure relative to another can be more challenging (e.g., amplitude of low frequency fluctuation [ALFF], voxel-mirrored homotopic connectivity [VMHC], ReHo). While the iFC similarity measure employed for initial gradient-based mapping efforts has several desirable features (e.g., ability to overcome noise through averaging across vertices, utility for full-brain parcellation, most homogeneous parcels), it is the only aspect of the intrinsic brain. It can be argued that studies may benefit from selection of their measures based upon the specific question or purpose at hand. For example, if the question is specifically about the transitions of fronto-parietal network, then dual-regression may be preferable. As might be expected, for DR-networks, the ICC of a given voxel showed profound dependencies on network membership and gradient strength; such properties should be considered in their application. Alternatively, more exploratory approaches may derive benefit from the calculation of multiple features for each cortical area simultaneously (e.g., fALFF, ReHo, VMHC, DC), providing gradient profiles which can be employed for analyses.

Among the functional indices examined in the present work, the gradients derived from ReHo (an index of local synchrony) were the most reliable, followed by similarity of iFC. Relative to other measures, these two measures are unique in that they involve averaged values, which will provide some degree of noise reduction. While the number of values being averaged into the ReHo score for a given vertex are dramatically fewer than those that go into the same vertex’s similarity of iFC, the values being averaged are inherently more similar to one another. Additionally, it is worth noting that ReHo is consistently noted to engender a high degree of reliability as a measure, which may in turn lend to its more reliable gradients (Chen et al. 2016; Jiang and Zuo 2015). It may be worth noting that ReHo relies on the non-parametric Kendall’s Concordance Coefficient rather than Pearson’s Correlation, which may also be a contributing factor (Zuo et al. 2013). Regarding similarity of iFC, two observations are worth noting. First, that the most closely related function indices were the dual regression components for the default and sensorimotor networks, which are most similar to the task-negative and task-positive networks defined by Fox et al. (2005). Second, those gradients found to be highly reliable for similarity of iFC, tended to have reliability across the other R-fMRI indices as well. These findings suggest summary measures (e.g., similarity of iFC) have clear strengths, though may have limitations with respect to their reliability to findings outside the cardinal large-scale networks (e.g., task-positive, task-negative).

### Limitations and Future Directions

Although promising, there are several limitations to the findings of the present work beyond those already discussed, which merit consideration. First, R-fMRI scans did not cover the whole brain for all the participants, particularly in the inferior temporal and orbital frontal cortex; as such, our ability to map transition profiles for these areas was limited. This reduced coverage could have impacted our findings (e.g. DR-auditory network), though we do not believe to a high degree. Second, while several studies have demonstrated that the areal transition profile based on R-fMRI correspond to the tasks activation (Wig et al. 2013; Laumann et al. 2015), the present work is limited in its ability to make generalizations regarding transition zone repeatability beyond R-fMRI; future serial scanning efforts would benefit from the inclusion of task fMRI as well. If the areal transition profile from task and rest are reliable, it may facilitate alignment across participants by offering spatial functional variability information to registration progress (Robinson et al. 2014). Finally, the current findings of reliable transition profile are based on datasets sampling a one-month period; future work may investigate the dynamic changes of areal organization across more extended periods of time, as well as address questions regarding potential age-related differences across the lifespan.

## Conclusions

In sum, the present work demonstrates the ability to map repeatable, individual-specific areal transition profiles for an array of iFC features, confirming their potential for fingerprinting and biomarker discovery. Data needs for achieving moderate to high test-retest reliability appeared to be greater than those for fingerprinting, though achievable. Points of convergence were noted in the cortical area transition zones defined using differing iFC indices. However, the distinctiveness of the full-brain transition zones profiles obtained using differing iFC indices suggests the merits of considering multiple indices to provide a more comprehensive characterization of cortical area transition zone properties.

## ACKNOWLEDGES

This study was supported by the Funds of International Exchange for Post-doctoral Scientists of China (2013), grants from the NIH (NIMH U01MH099059 to M.P.M.; NIMH BRAINS R01-MH101555 to R.C.C.), the Natural Science Foundation of China (81270023, 81278412, 81171409, 81000583, 81471740, 81220108014), the National Science Foundation for Post-doctoral Scientists of China (No. 2013M530073), the Key Research Program and the Hundred Talents Program of the Chinese Academy of Sciences (KSZD-EW-TZ-002, X.N.Z), as well as gifts from Phyllis Green, Randolph Cowen and Joseph Healey to M.P.M‥ All of the authors declare no conflict of interest. We would also like to thank Ian Hickie, Katie McMahon, Paul Thompson, and Greig de Zubicaray, who are investigators for the QTIM project and generously provided the data (Australian Twin Study; R01HD050735). We also thank Evan Gordon and Timothy Laumann for valuable discussions on surface-based gradient computation.

---

**Figure S1.**
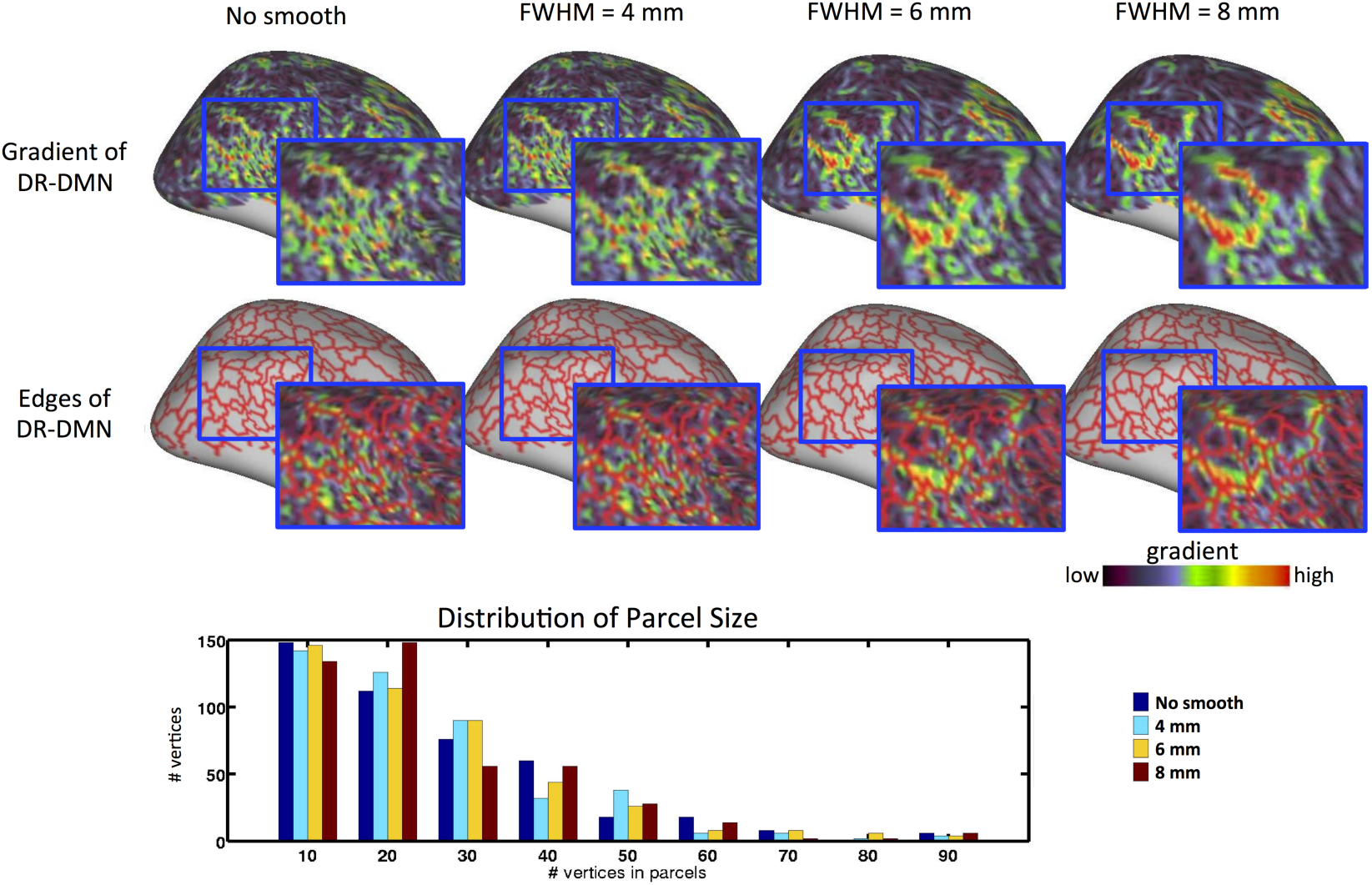
The effect of smooth filtering kernel size to the gradient. The larger the smoothing radius is, the lower the sensitivity to noise, resulting larger parcel size. In the present study, given that the resolution of final ‘fsaverage5’ surface is ~3.5 mm (distance between one-step neighbors of middle surface of ‘fsaverage5’), we employed FWHM=8 mm (about twice of the neighbors distance) smoothing to the map of functional indices (iFC similarity, ReHo, DC, EC, DR-Networks), to ensure the FWHM covers one-step neighbors for all the vertices on the surface mesh.

**figure S2.**
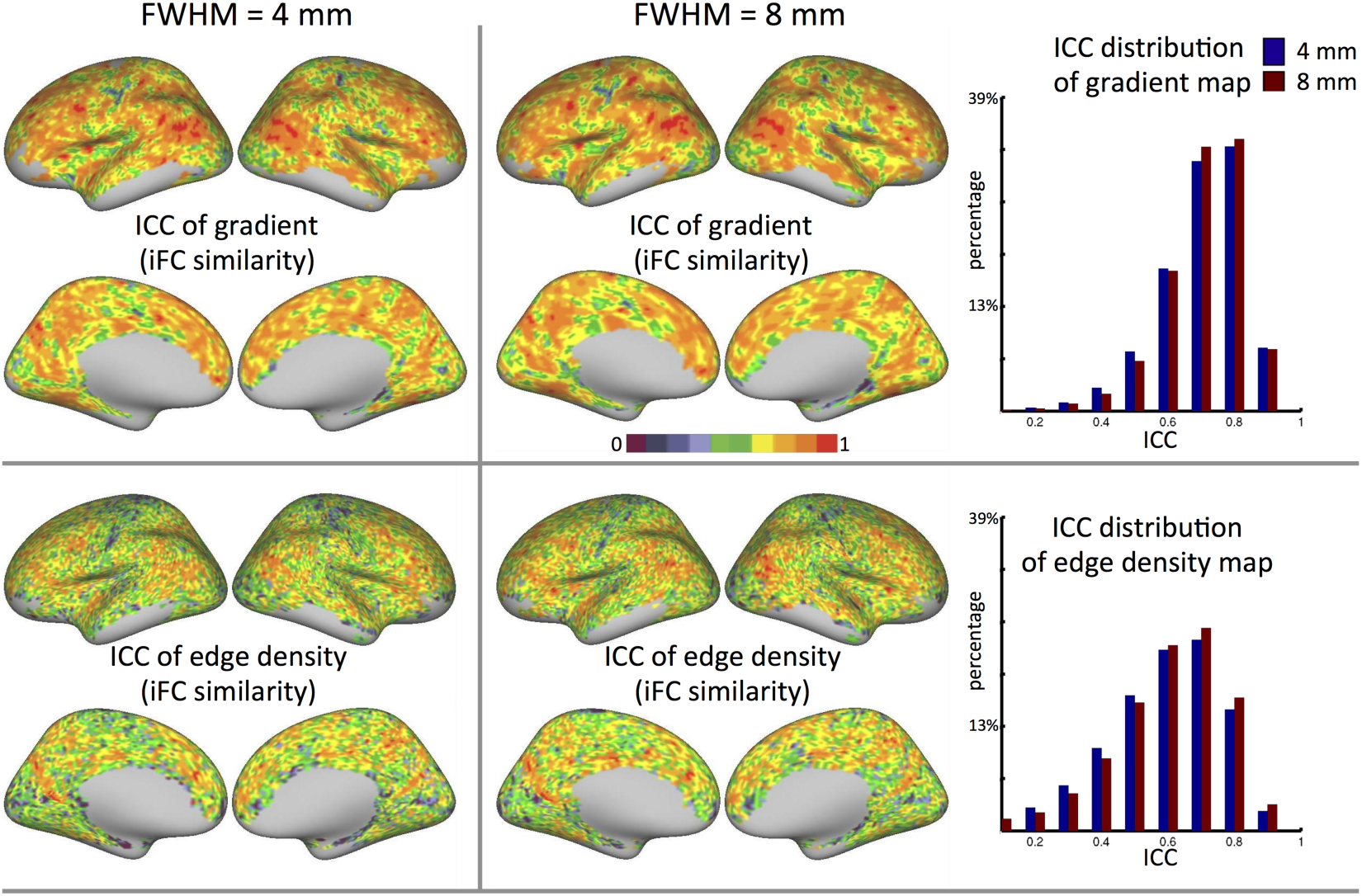
The effect of smooth filtering kernel size to the ICC. Inherent to this procedure is a smoothing step performed before edge detection to reduce noise as well as spatial variance of gradients. ICC obtained from larger smoothing parameter (FWHM=8) was slightly higher than small smooth filter (FWHM=4).

**figure S3.**
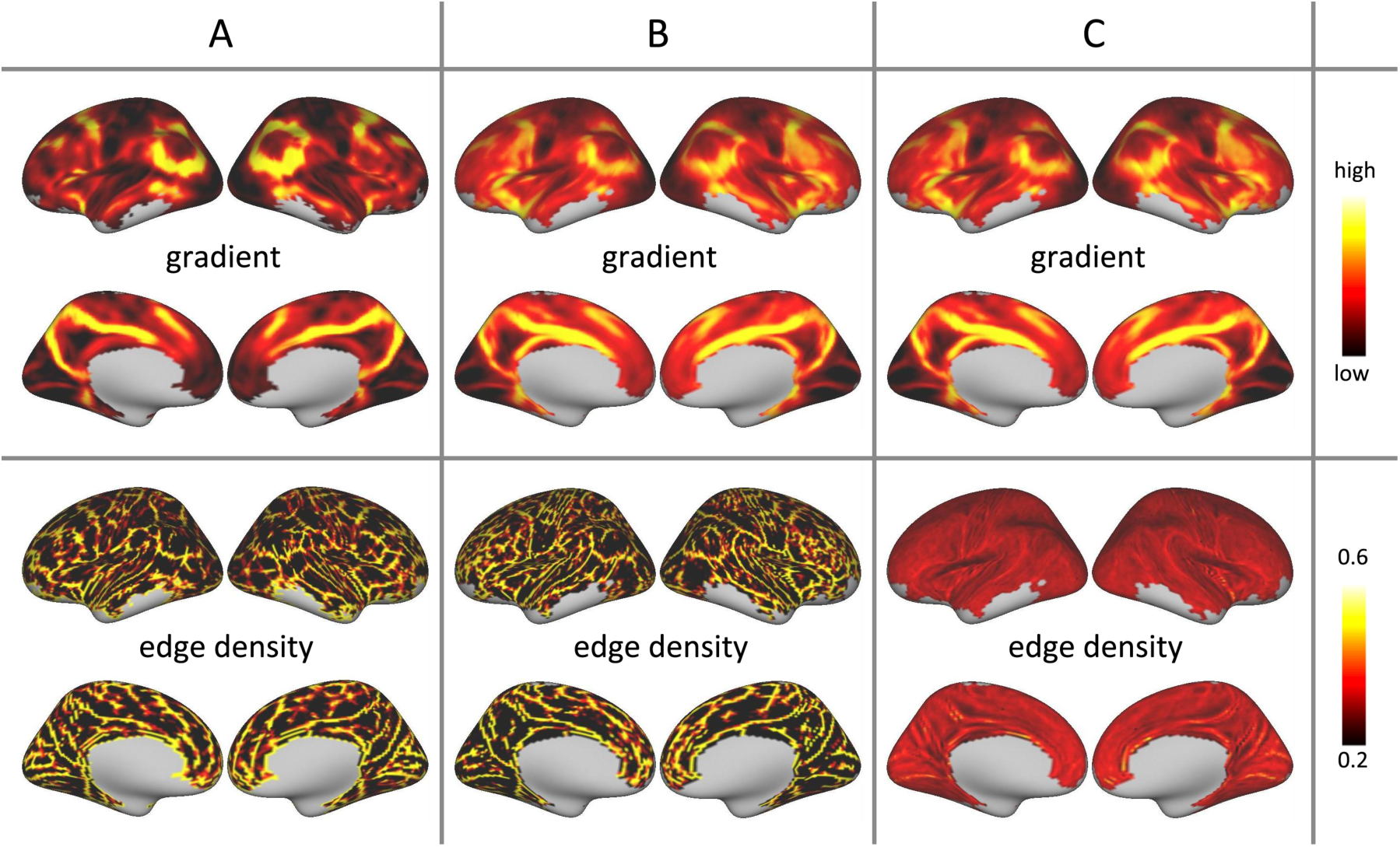
Group-level gradient and edge density maps of iFC similarity of three different pipelines based on 400 datasets (de Zubicaray et al., 2008). A: The iFC similarity matrixes were averaged across participants first to yield group iFC similarity matrix and then calculated group gradients and edge density on *‘fsaverage5’* template surface (Wig et al., 2013). B: The gradients were calculated on ‘fsaverage5’ version of native surface at individual level and then averaged across participants to yield group gradient map followed by edge detection on template surface (Gordon et al., 2014). C: The same pipeline applied in the present study that iFC similarity, gradients and edge density are all computed on ‘fsaverage5’ version of native surface at individual level and then averaged across participants for the final group map. Of note, to keep the analysis comparable to previous study, the same non-maxima suppression procedure (Wig et al., 2013) was employed to detect edges for those three pipelines, which labels a vertex as an edge if its gradient is larger than the gradients of at least two non-adjacent pairs of its neighbors.

**figure S4.**
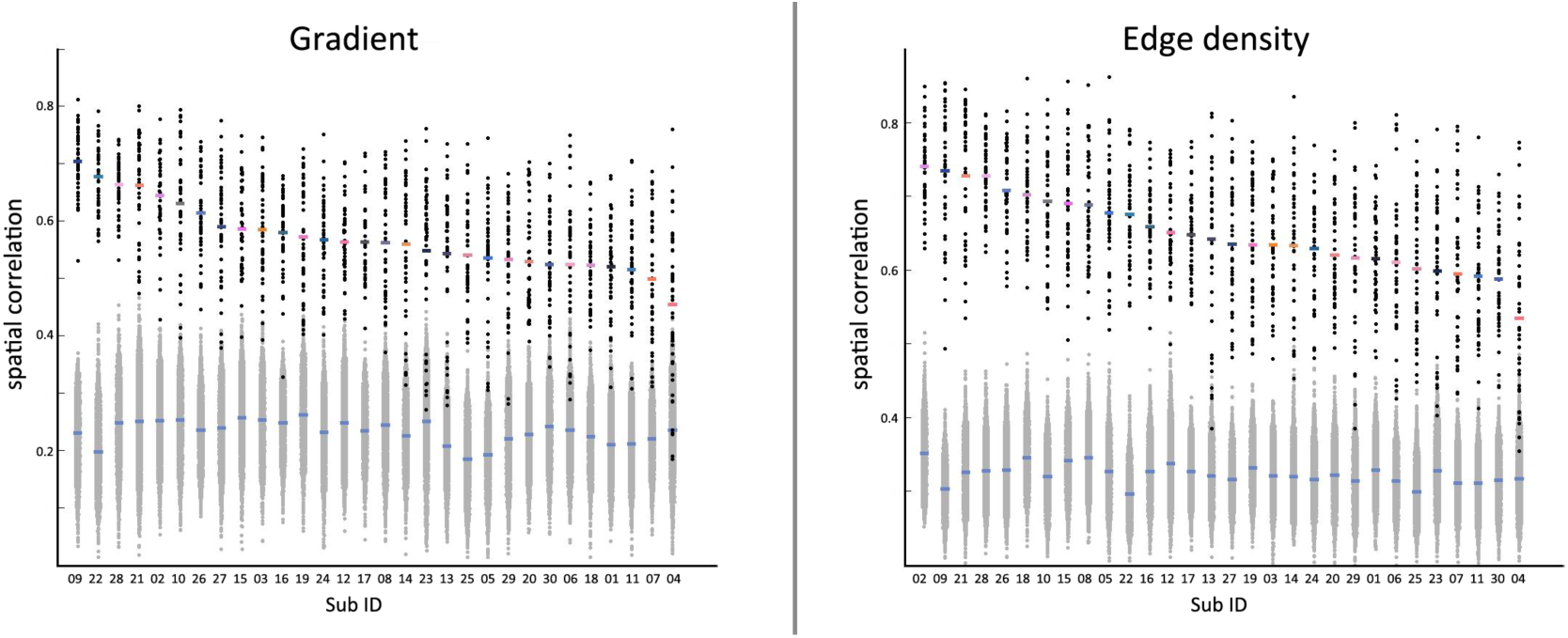
The distribution of spatial correlations for each participant (ordered by mean of the within-sindividual correlations). Black dots represent the correlations of gradient (left panel) and edge density (right panel) between any two 10-minute sessions within the same participant. Grey dots depict the correlations of gradient between any two 10-minute sessions from different participants. The mean or standard deviation of within-individual correlations are not associated with age, gender, head motion or functional-to-structural co-registration.

**figure S5.**
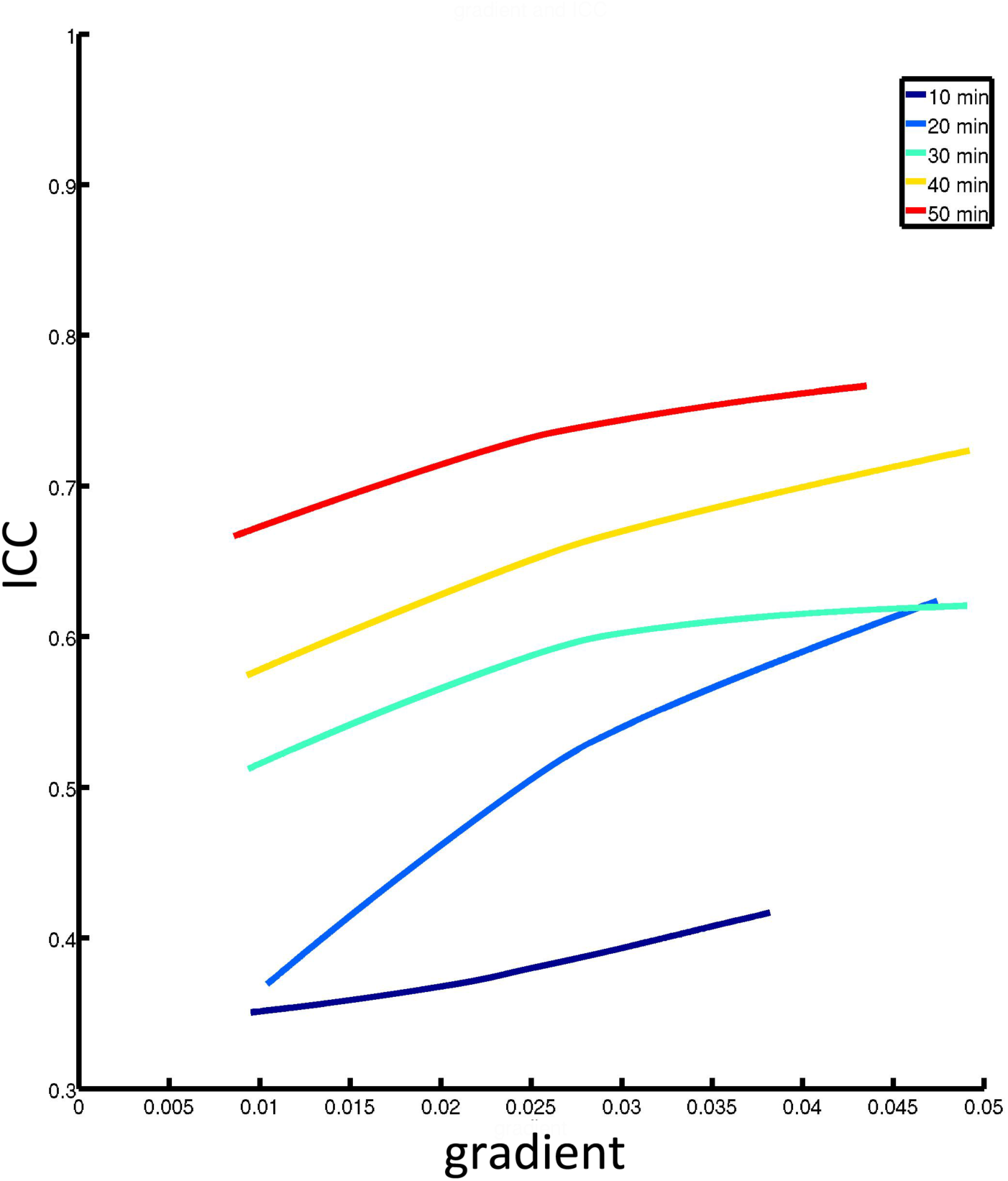
The distribution of ICC is slightly related to gradient scores of iFC similarity. Each line represents the association (LOWESS fit) between ICC with gradient strength for each of the scan durations of 10min, 20min, 30 min, 40 min, and 50 min.

**figure S6.**
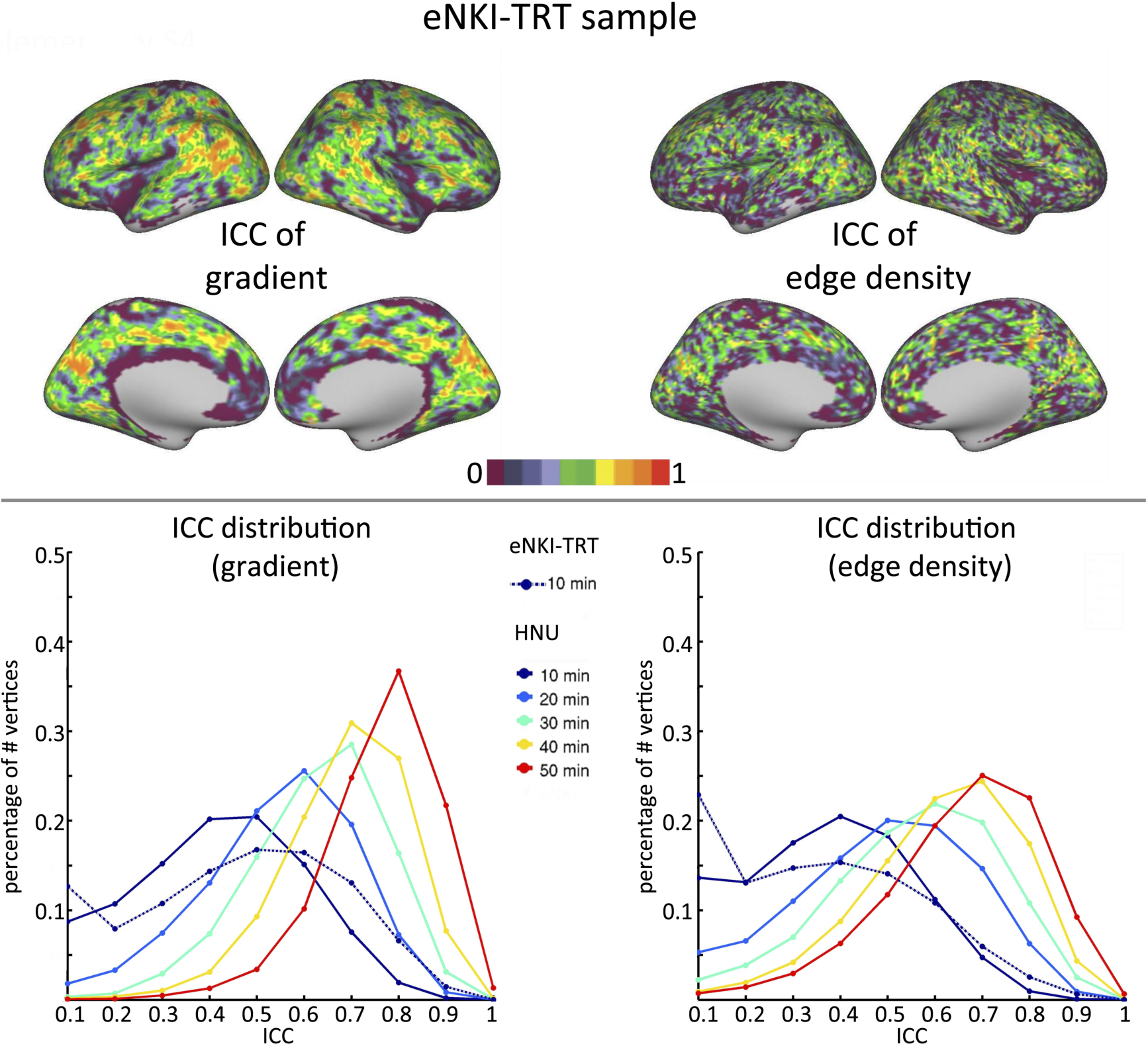
The ICC maps for the gradient and edge density of iFC similarity (top panel) based upon eNKI-TRT sample (TR=645 ms, 10 min). The histograms of ICC distribution are plotted with dash line at the bottom as well as the ICC distribution from HNU dataset (TR=2000 ms, 10 min, 20 min, 30 min, 40 min, 50 min).

**figure S7.**
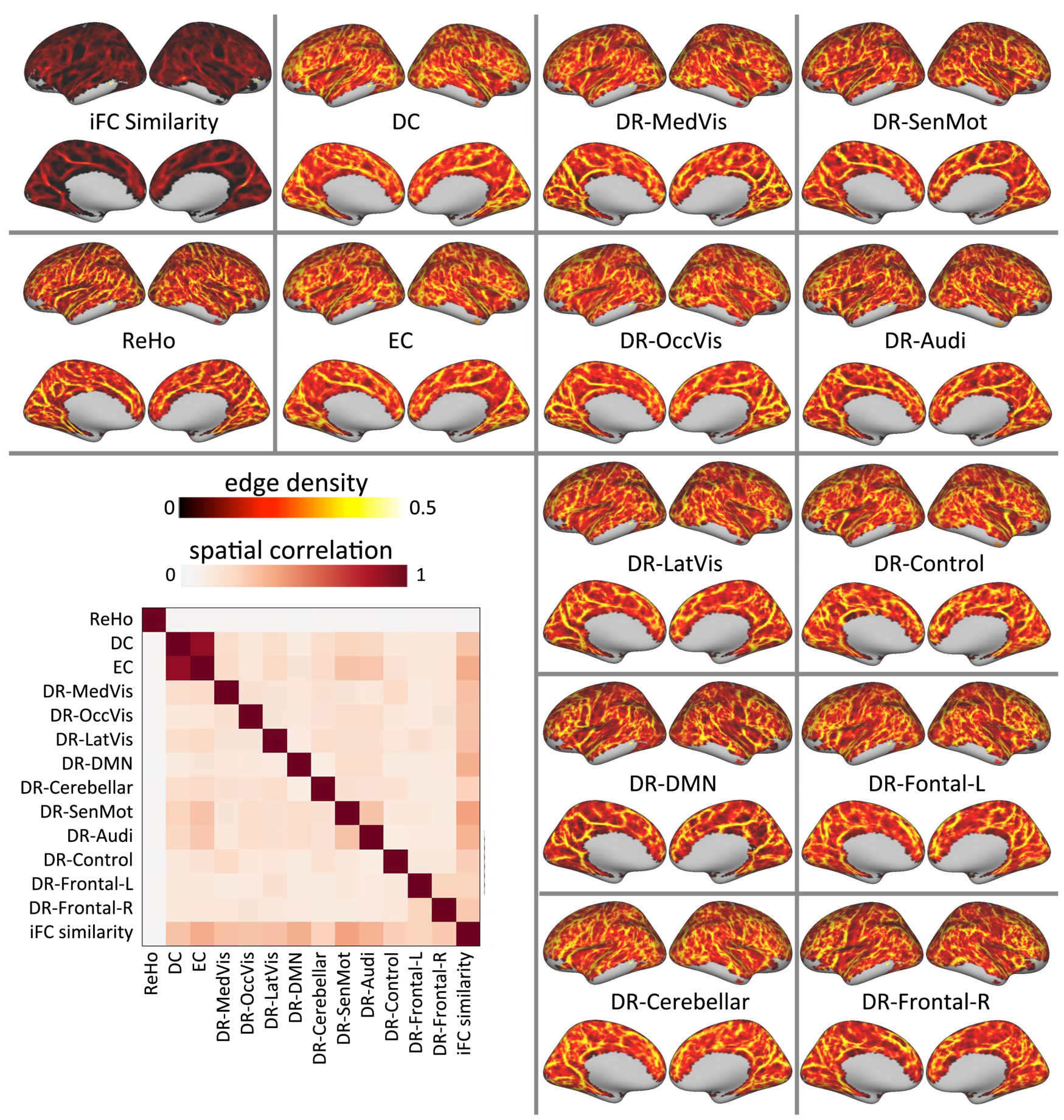
Group-level edge density maps for each functional indices from two 50-minutes subsets. For each participant, we calculated the spatial correlation matrix of the edge density maps for various functional indices, and then averaged across individuals to provide the spatial correlation matrix (bottom left corner).

**figure S8.**
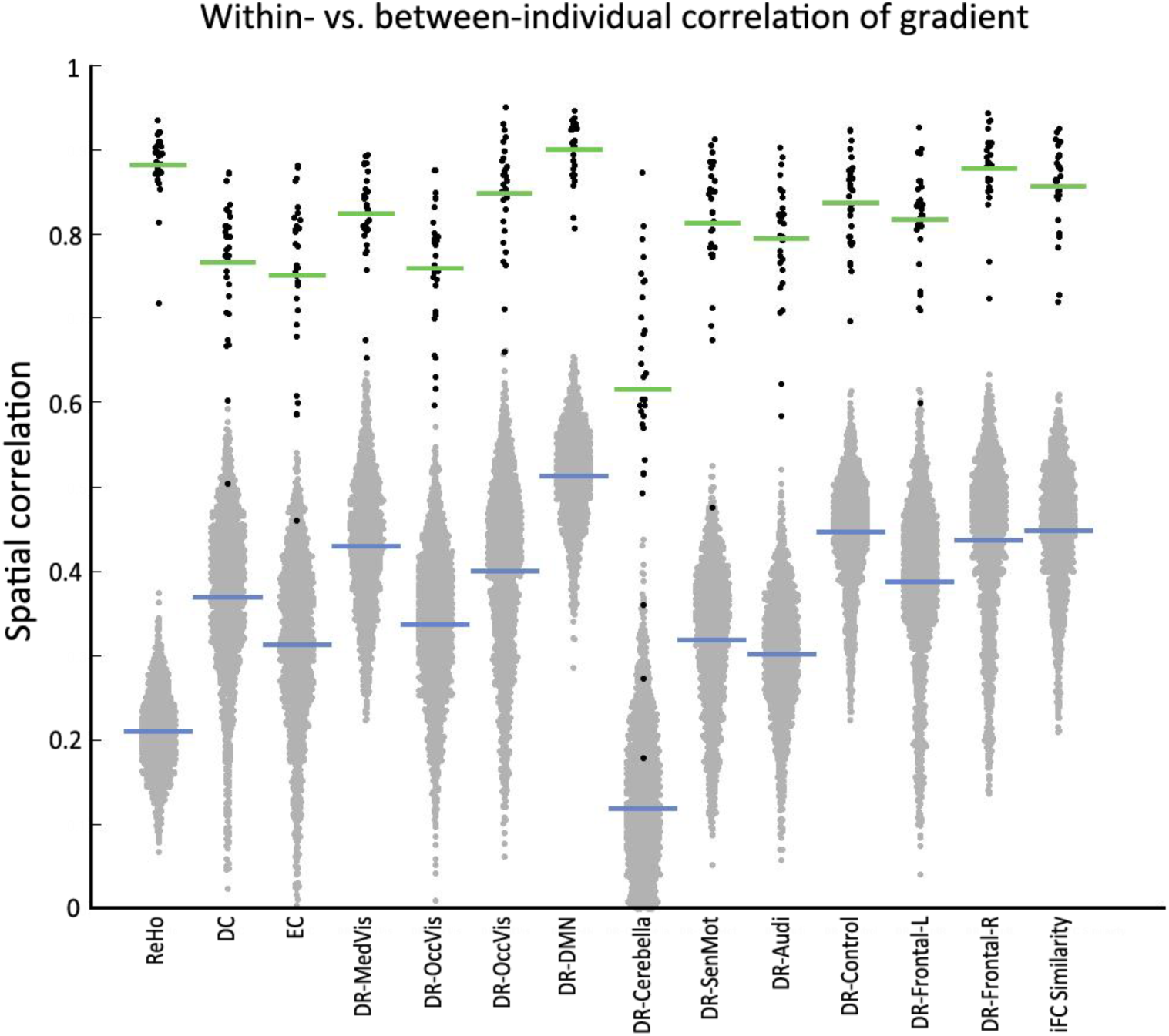
The distribution of within- versus between-individuals correlation for the gradients of 14 functional indices. The Black dots are within individual correlations between two 50-min subsets for each individual while grey dots are correlations between two 50-min subsets from different individuals.

**figure S9.**
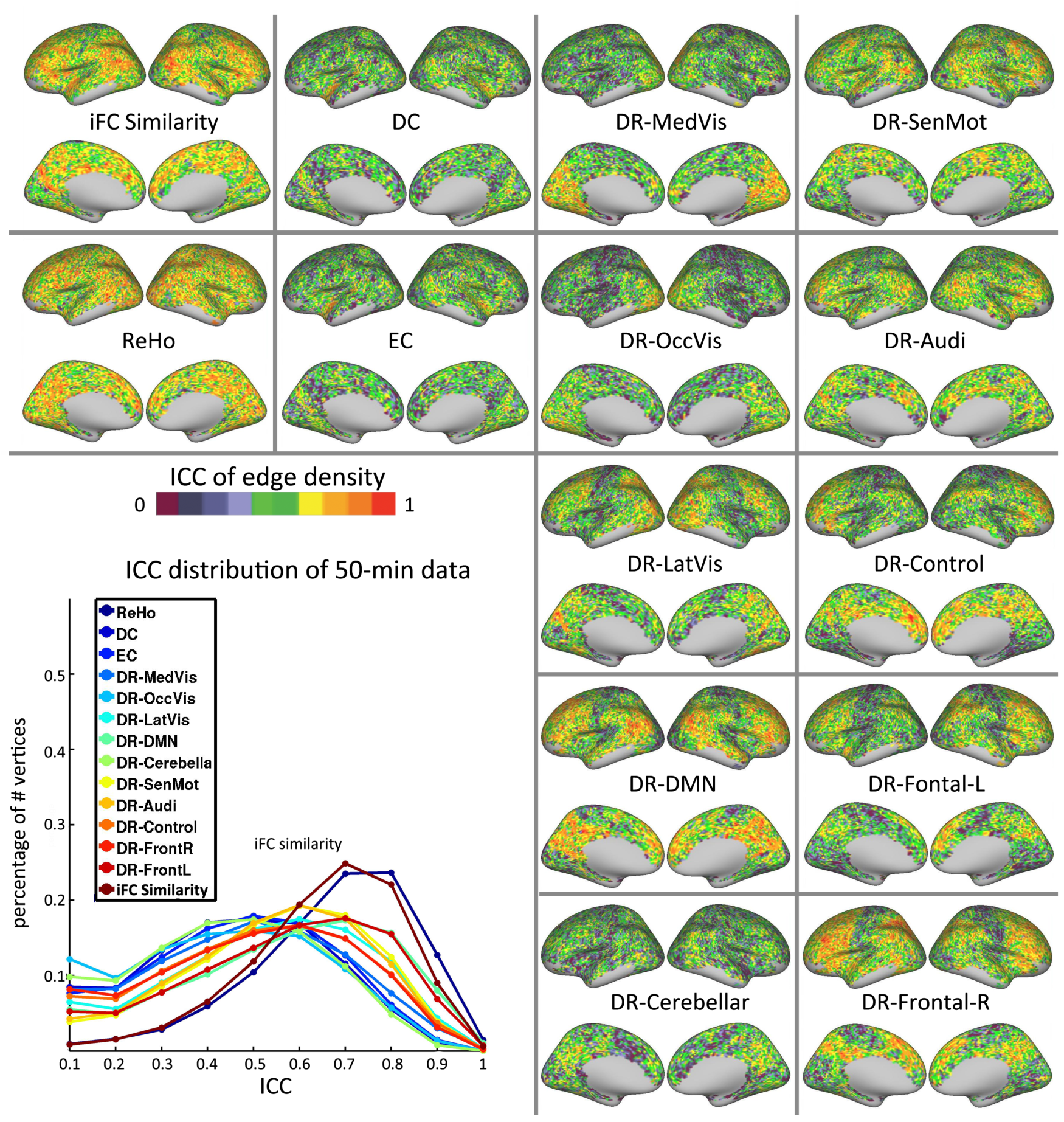
The intra-class correlation maps for the edge density of 14 functional indices are depicted here (ICC was calculated between two 50-minute subsets of data generated through random selection [without replacement]). The distribution of ICC for edge density maps for each functional metric is shown in the bottom-left corner.

**figure S10.**
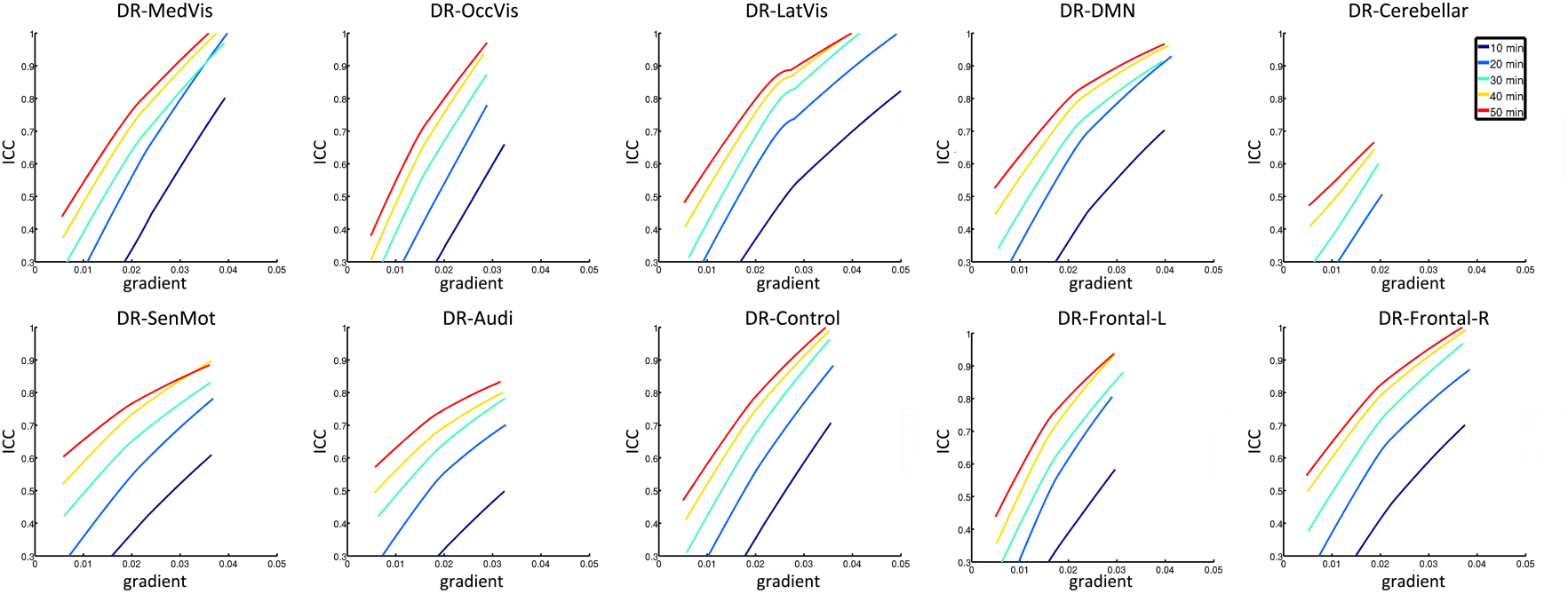
The relationships between ICC and gradient strength across vertices for DR-networks. Each line (LOWESS fit) represents the association of ICC with gradient strength for each of the scan durations of 10min, 20min, 30 min, 40 min, and 50 min.

**figure S11.**
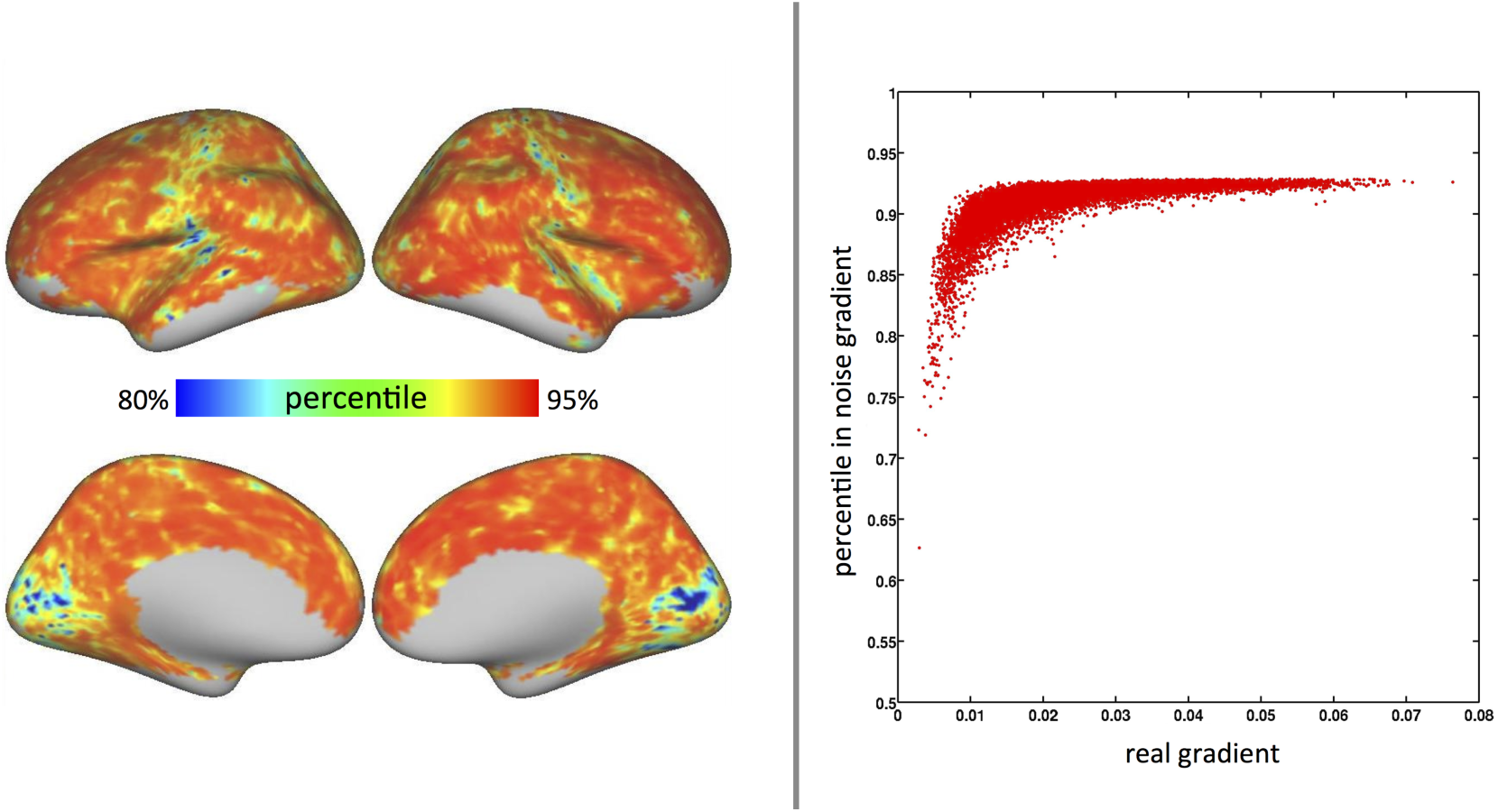
The percentile of gradients from real data in the frequency distribution of gradients from random noise for a representative participant (left). The scatter plot (right) shows that where the lower percentile of real gradients, a relatively lower gradient score in the real data.

## Footnotes

1 We made use of the Yeo et al. 2011 network definitions rather than those of Smith et al., 2009 networks due to: a) the lack of confidence maps, and b) overlaps that occur among the network maps when transformed into surface space.!

